# Combining SIMS and mechanistic modelling to reveal nutrient kinetics in an algal-bacterial mutualism

**DOI:** 10.1101/855999

**Authors:** Hannah Laeverenz Schlogelhofer, François J. Peaudecerf, Freddy Bunbury, Martin J. Whitehouse, Rachel A. Foster, Alison G. Smith, Ottavio A. Croze

**Affiliations:** Cavendish Laboratory, University of Cambridge, United Kingdom; Institute of Environmental Engineering, Department of Civil, Environmental and Geomatic Engineering, ETH Zürich, Switzerland; Department of Plant Sciences, University of Cambridge, United Kingdom; Swedish Museum of Natural History, Stockholm, Sweden; Department of Ecology, Environment and Plant Sciences, Stockholm University, Sweden

## Abstract

Microbial communities are of considerable significance for biogeochemical processes, for the health of both animals and plants, and for biotechnological purposes. A key feature of the interactions between microbes is the exchange of nutrients between cells. Isotope labelling followed by analysis with secondary ion mass spectrometry (SIMS) can identify nutrient fluxes and heterogeneity of substrate utilisation on a single cell level. Here we present a novel approach that combines SIMS with a mechanistic model to reveal otherwise inaccessible nutrient kinetics. The method is applied to study the onset of a synthetic mutualistic partnership between a vitamin B_12_-dependent mutant of the alga *Chlamydomonas reinhardtii* and the B_12_-producing, heterotrophic bacterium *Mesorhizobium loti*, which is supported by algal photosynthesis. Results show that an initial pool of fixed carbon delays the onset of mutualistic cross-feeding, and the model allows quantification of this delay. Our method is widely applicable to other microbial systems, and will contribute to furthering a mechanistic understanding of microbial interactions.

## Introduction

Microbial communities underpin many globally important processes, from biogeochemical cycles (1) and the ecology of aquatic (2, 3) and terrestrial food webs (4, 5), to wastewater treatment (6, 7) and the health of agricultural soils (8). A key feature of the interactions within these communities is the exchange of metabolites between species (9). In aquatic environments, photosynthetic carbon fixation by phytoplankton supports higher trophic levels, but also provides an important carbon source for heterotrophic bacteria (10–12). Conversely, bacteria have been shown to provide limiting nutrients to algae, including nitrates, phosphates and iron (13), vitamins (14, 15) and carbon dioxide (16). Depending on environmental conditions, these metabolite exchanges control the outcome of microbial interactions, from parasitic, through commensal, to mutualistic (17–19).

To exploit microbial communities for biotechnological applications, it is crucial to be able to predict and control microbial interactions. Extensive studies of natural microbial communities using metagenomics, metatranscriptomics and metaproteomics have provided considerable insight into potential metabolite exchanges (20, 21). However, to obtain a fully predictive, mechanistic understanding of microbial interactions it is also essential to use bottom-up approaches employing laboratory model systems and mathematical models (22–25). For example, the comparison of a nutrient-implicit Lotka-Volterra model with experiments studying co-cultures of genetically engineered strains of yeast that each provide a different essential nutrient to the other demonstrated a limiting nutrient-induced shift from mutualism via parasitism to competition (26). Moreover, studies of engineered yeast communities combining agar pad experiments and models incorporating nutrient diffusion revealed that cross-feeding interactions influence genetic drift during spatial expansion (27), and that spatial self-organisation favours cooperation over cheating (28).

The exact metabolic interactions within microbial communities are often unknown. Secondary ion mass spectrometry (SIMS, NanoSIMS), an imaging mass spectrometry technique capable of analysing single microbial cells, reviewed in (29–33), has been instrumental in identifying new symbioses and microbial interactions for both cultured and non-cultured associations (34–37). Moreover, the metabolic activity and phylogenetic identity (16S rRNA) of single cells can be linked by combining *in situ* hybridization methods with SIMS (38, 39). Using SIMS and NanoSIMS to visualise and quantify substrate utilisation in single cells, filaments, and colonies of microbial cells has helped to determine the heterogeneity of single cell metabolic activity (38, 40), sub-cellular location of assimilated substrates (41, 42), nutrient exchanges between symbiotic partners (35, 36) and the effect of physical attachment on carbon and nitrogen fluxes between bacteria and microalgae (43, 44).

In these studies, SIMS was primarily used to visualise and measure nutrient assimilation and transfer. In the dilute aquatic environment, microbial interactions will involve dynamic nutrient exchanges, particularly at the onset of association, when metabolite fluxes may be quite different from those arising during a stable, long-term interaction. Here we explore the establishment of mutualistic interactions with a well-characterised model system: a co-culture of the cobalamin (vitamin B_12_) dependent, photosynthetic alga *Chlamydomonas reinhardtii metE7* strain (45) and the B_12_-producing, heterotrophic bacterium *Mesorhizobium loti*. Previous studies of this system, and a closely related one comprising the naturally B_12_-dependent alga *Lobomonas rostrata*, have demonstrated mutualistic growth dynamics predicated on the exchange of vitamin B_12_ and organic carbon photosynthate (45, 46). The relative proportions of the two organisms are stably maintained over hundreds of generations, but can be perturbed by supplementation with cobalamin or an organic carbon source like glycerol (46). The effect of environment geometry on the mutualistic dynamics of spatially separated populations was also recently modelled mathematically, and realised experimentally (47). Here, SIMS experiments that follow the temporal variation in ^13^*C* labelling are combined with a mechanistic, nutrient-explicit model to gain further insight into how these organisms interact. The model permits use of the SIMS data to obtain nutrient exchange kinetics, which were not possible to measure experimentally, and to explore potential mechanisms for the observed single cell heterogeneity.

## Materials and Methods

### Algal and bacterial strains

The B_12_-dependent alga used in this work was *C. reinhardtii metE7* (ref. 45). The B_12_-producing bacterium used was *M. loti* (MAFF 303099), originally a gift from Prof Allan Downie, John Innes Centre, UK.

### Growth conditions

All cultures were grown in a 12 *h* − 12 *h* light-dark cycle at 25°*C*, shaking at 120 *rpm*. The light intensity of the photosynthetically active radiation was approximately 70 μ*mol m*^−2^ *s*^−1^, measured using a Skye PAR sensor (SKP 215). Tris-minimal medium was used for all cultures, meaning that *C. reinhardtii metE7* grew phototrophically in our experiments. Tris-minimal medium is based on TAP (48) but omits the acetic acid and *HCl* is used to titrate to *pH* 7 (ref. 49). The trace elements solutions used (Supplementary Table S1) were adapted from (50) to include a seventh solution containing cobalt, since cobalt is required as the central ion of vitamin B_12_. The cobalt concentration was chosen to be the same as in Hutner’s trace elements (51). Cyanocobalamin (referred to as B_12_ throughout this work), glycerol and sodium bicarbonate were added to the medium as required (Supplementary Table S2).

Dissolved sodium ^13^*C*-bicarbonate (Sigma-Aldrich *NaH*^13^*CO*_3_, 98 *atm*% ^13^*C*) was used for the stable isotope labelling of microbial cultures (the work-flow is illustrated in Supplementary Figure S1). A sample taken from the 600 *mL* axenic pre-culture of algae was washed and then resuspended in 1 *L* of fresh media containing 100 *ng L*^−1^ B_12_ and 5 *mM NaH*^13^*CO*_3_. This pre-labelling culture of algae was grown for 48 *h* (see Supplementary Information for the experimental and model results for this culture). An axenic pre-culture of bacteria was grown in media with 0.1 % (*v*/*v*) glycerol, which was then sampled, washed and re-suspended in 750 *mL* fresh media containing 5 *mM NaH*^13^*CO*_3_, to which 250 *mL* of pre-labelled algae was added to initiate the co-culture. Cultures of axenic bacteria were grown with 5 *mM NaH*^13^*CO*_3_ and different concentrations of unlabelled glycerol.

### Population growth

Population growth was monitored using viable counts. A series of 10-fold dilutions were performed and aliquots of 20 μ*L* from relevant dilutions (i.e. chosen such that approximately 10 to 100 colonies would result after plating) were spotted onto TY agar plates. The plates were tilted back and forth to disperse the cells and make the colonies easier to distinguish (52). Plates were incubated in continuous light at 25°*C* for approximately 5 days and in the dark at 30°*C* for approximately 2 days, for algal and bacterial colonies respectively. Two independent viable counts were obtained for each time-point and the results converted to values for the population size in units of colony forming units per unit volume (*cfu mL*^−1^).

### Isotope Ratio Mass Spectrometry

Isotope Ratio Mass Spectrometry (IRMS) was used to measure ^13^*C* ratios for bulk samples of algal and bacterial biomass. IRMS also measured the total carbon and nitrogen content, which was used to calculate the C:N ratio and, together with dry mass and cell density measurements, to estimate the carbon yield (i.e. *cells molC*^−1^) for algae and bacteria, see Table S4 and Supplementary Information for details.

### Secondary Ion Mass Spectrometry

#### Sample preparation

Below is a brief outline of the SIMS sample preparation procedure, full details are in Supplementary Information. Samples were chemically fixed using formaldehyde. Vacuum filtration was used to deposit the cells onto 0.22 μ*m* pore size membrane filters with a ≈ 20 *nm* gold coating, with nucleic acid staining and confocal microscopy (Olympus Fluoview FV1200) used to confirm an even distribution of cells on the filter. A single hole punch was used to cut out 4 − 6 *mm* disks from the filter samples. Following this, a Zeiss laser micro-dissection microscope (Zeiss LSM710-NLO housed at the LCI facility of the Karolinska Institute, Stockholm) was used to image the autofluorescence of the algal chlorophyll and create laser marks on the samples, used to locate areas of interest with the camera of the SIMS instrument. Lastly, the samples were placed on conductive sticky tape, mounted onto a glass disk and sputter coated with gold.

#### SIMS analysis

SIMS analysis was performed using the Cameca IMS 1280 at the NordSIM facility in the Department of Geosciences at the Swedish Museum of Natural History in Stockholm. The instrument uses a Gaussian focussed primary ion beam of caesium ions (*Cs*^+^). For selected positions on the filter sample, 45 × 45 μ*m* square areas were pre-sputtered for 10 *s* with a 3 *nA* primary ion beam. Within this pre-sputtered region, 100 scans of a 35 × 35 μ*m* square area were measured using a ≈ 60 − 80 *pA* primary ion beam (spot size of approximately 1 μ*m*). The secondary ion mass peaks were measured using an ion counting electron multiplier in peak hopping mode with a 44 *ns* electronically gated dead-time. The count times for the ^12^*C*^14^*N*^−^, ^12^*C*^15^*N*^−^ (not used in subsequent analysis) and ^13^*C*^14^*N*^−^ secondary ion peaks were 1, 0.5 and 2 *s* respectively. A mass resolution (*M*/Δ*M*) of approximately 6000 for the preliminary experiments (see Figure S10) and 7000 for the final experiments was used; a mass resolution of 6000 − 7000 was sufficient in resolving both the ^12^*C*^14^*N*^−^ and ^13^*C*^14^*N*^−^ peaks. Interference of ^11^*B*^16^*O*^−^ with the ^13^*C*^14^*N*^−^ peak at mass 27 was not an issue because no boron or boron containing compounds were used in the culture media. The SIMS measurements were run once for bacterial cells and repeated 1 − 8 times for each algal cell. The WinImage2 software (Cameca) was used to obtain the isotope ratio *R* = ^13^*C*/^12^*C* for single cells of algae and bacteria (see Supplementary Information for details). The atomic fraction of ^13^*C*, i.e. *f* = ^13^*C*/(^13^*C* + ^12^*C*), was calculated using

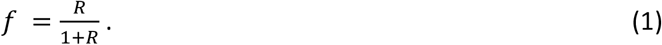

Several technical considerations were taken into account (full details in Supplementary Information and Figure S2). First, a depth analysis was performed by taking repeated measurements of the same cells, which demonstrated that a single measurement was sufficiently representative for bacteria, whereas for algal cells the mean of three repeated measurements was used to obtain a representative measurement. Second, a scattering effect associated with highly labelled algae was observed, therefore for the analysis described in this work only bacteria from scan areas not containing labelled algae were included. Lastly, the dilution effect, due to chemical fixation and nucleic acid staining introducing unlabelled carbon into cells, was taken into consideration (see Table S3). To estimate the undiluted atomic fraction of ^13^*C*, SIMS results were *dilution-corrected* using the method established in (53).

### Mechanistic model

To better understand the carbon kinetics revealed by the co-culture experiments and the underlying mutualistic microbial dynamics, a mechanistic model was formulated. A brief overview is provided here with full details given in Supplementary Information. The model captures essential nutrient exchanges between algae and bacteria, shown schematically in Figure 1. Algal growth is dependent on the external B_12_ concentration *v*, originating from bacterial production. The growth of bacteria, in turn, depends on the external concentration of algal-derived DOC, modelled as an effective single carbon source *c*_*o*_. This exchange of B_12_ and DOC provides mutualistic coupling between the two species. The co-culture is assumed to be well-mixed, such that

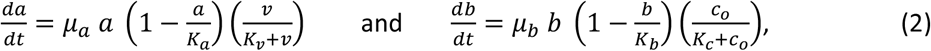

with *a* and *b* the algal and bacterial cell densities respectively, *μ*_*a*_ and *μ*_*b*_ the maximum growth rates, *K*_*a*_ and *K*_*b*_ the carrying capacities, and *K*_*v*_ and *K*_*c*_ the half-saturation concentrations. Although DIC is assumed to be non-limiting (as in the experiments), accounting for DIC kinetics was essential to connect the model to SIMS experiments, where isotope labelling relied on assimilation of ^13^*C* via DIC. As any living cell, heterotrophic bacteria can assimilate inorganic carbon through carboxylation reactions (54, 55). The model incorporates this observation through a DIC uptake parameter defined as 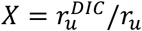, where 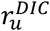 is the DIC uptake rate and *r*_*u*_ the total carbon uptake rate. Bacterial respiration further contributes to the inorganic carbon kinetics (56). This is modelled through the maximum bacterial growth efficiency *η*, which quantifies how respiration affects carbon uptake in the exponential growth phase. For *η* → 1, respiration goes to zero and does not affect carbon uptake. Instead, with *η* → 0 respiration rate is high compared to growth rate and thus strongly affects the carbon kinetics. Further, the model minimally describes photosynthesis and carbon storage in algae by splitting algal carbon biomass into two internal components, *photosynthetically-active* carbon *c*_*a,p*_, available for exudation, and *stored* carbon *c*_*a,s*_, used for biomass growth, in storage compounds (e.g. starch) and for cellular maintenance. Thus, the model effectively describes DOC exudation as originating from excess algal photosynthesis. The vitamin, DOC and DIC concentrations in the model are governed by the rate laws

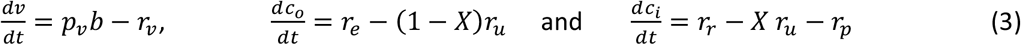

respectively, where we assume a constant B_12_ production rate per bacterial cell *p*_*v*_, *r*_*v*_ is the total vitamin uptake rate by algae, *r*_*e*_ the total DOC exudation rate by algae, *r*_*u*_ the total carbon uptake rate by bacteria, *r*_*r*_ the total bacterial respiration rate and *r*_*p*_ the total photosynthetic carbon assimilation rate by algae. The DOC exudation rate from algae is given by *r*_*e*_ = (1 − *ϕ*_*s*_) *p*_*c*_ *a*, where *p*_*c*_ determines the DOC production rate per algal cell (assumed constant), and *ϕ*_*s*_ defines the fraction of carbon *stored* by algae, i.e. *ϕ*_*s*_ = *c*_*a,s*_/(*c*_*a,s*_ + *c*_*a,p*_). Combining the differential equations for the carbon concentrations and the definition of atomic fraction *f* = ^13^*C*/(^13^*C* + ^12^*C*), we can write down differential equations for the dynamics of the atomic fractions, observed experimentally using SIMS. As an example, the atomic fraction for bacteria is given by

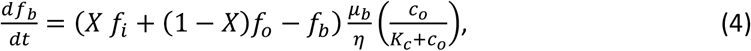

with *f*_*b*_, *f*_*i*_ and *f*_*o*_ the atomic fractions of ^13^*C* for bacteria, DIC and DOC respectively, and all other parameters as previously defined.

**Figure 1:**
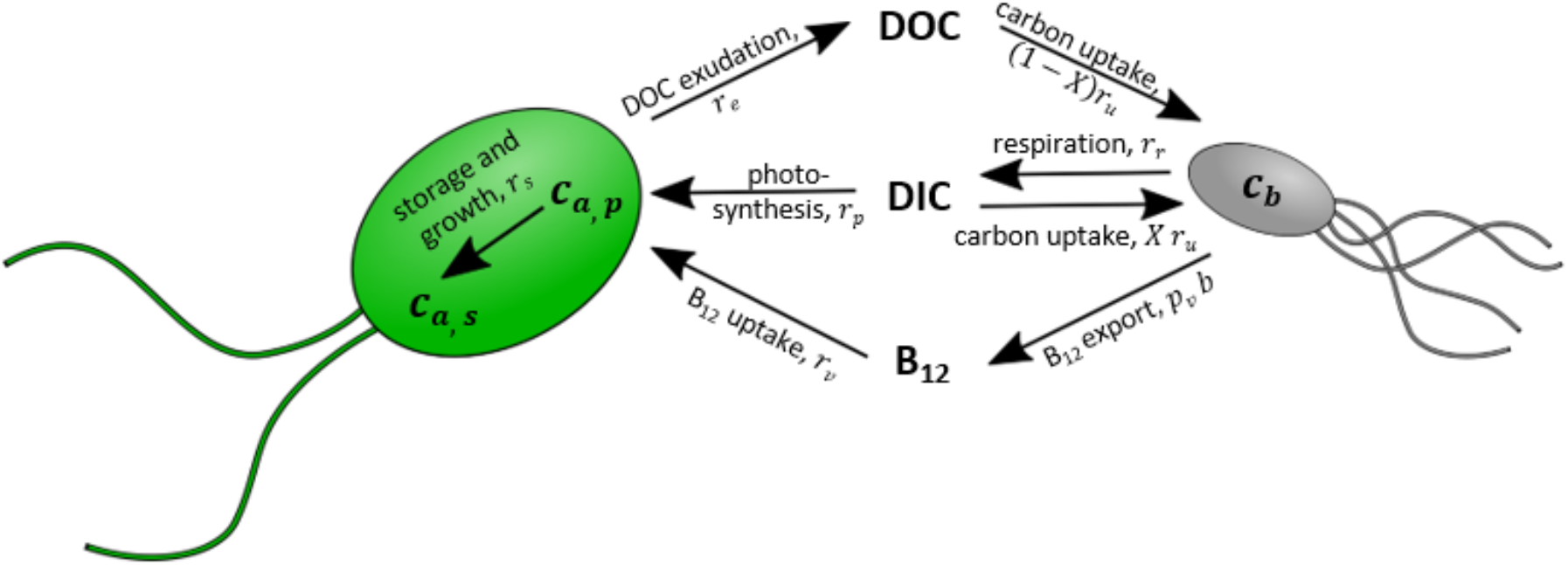
Schematic to illustrate the nutrient kinetics included in the algal-bacterial co-culture model. Vitamin B_12_ is released by bacteria and required for algal growth. Bacterial growth is dependent on DOC produced by algae. Also considered are: algal photosynthesis, carbon storage, and DOC exudation from excess photosynthesis; bacterial respiration and DIC uptake. An overview of the model is given in Materials and Methods with full details provided in Supplementary Information.

## Results

### Inorganic carbon acquisition by axenic bacteria

Axenic cultures of the rhizobial bacterium *M. loti* provided a benchmark for applying our method to the co-culture and allowed quantification of bacterial inorganic carbon acquisition. *M. loti* was grown axenically for 72 *h* with 5 *mM NaH*^13^*CO*_3_ (the labelled DIC source) and different concentrations of unlabelled glycerol, providing a source of organic carbon. SIMS images (Figure 2A) were used to determine the atomic fraction of ^13^*C*, *f*, for individual bacterial cells. The quantity *f*_*b*_ (Figure 2B) represents the fraction of ^13^*C* averaged over a distribution of single cell measurements; single cell heterogeneity effects are discussed below.

**Figure 2:**
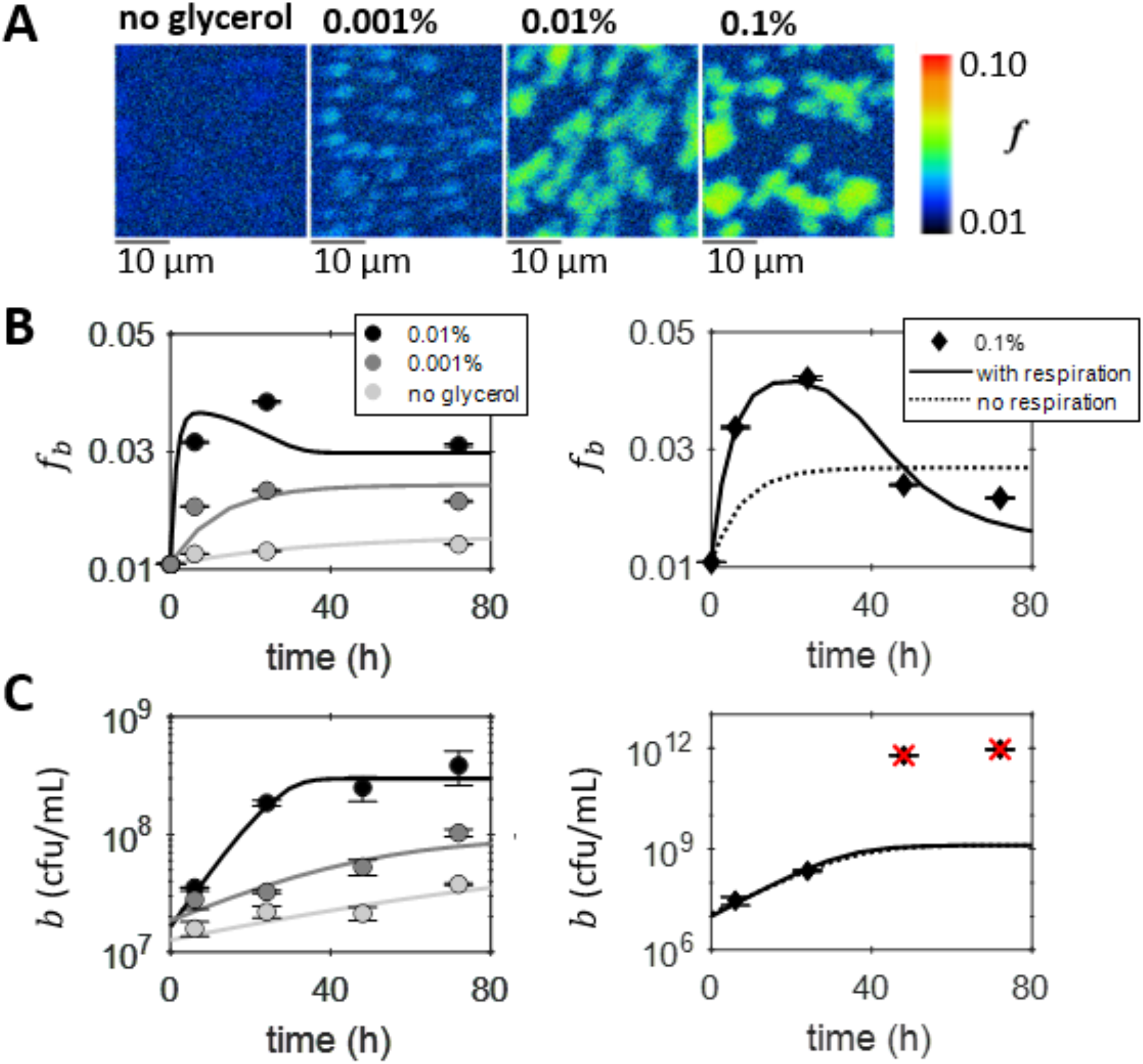
Inorganic carbon acquisition by axenic bacteria. (A) Example images of the atomic fraction of ^13^*C*, *f*, obtained by SIMS analysis of bacterial cells sampled after 24 *h* of axenic cultures grown with different concentrations of unlabelled glycerol and 5 *mM NaH*^13^*CO*_3_. The colour map shows the scale, starting at natural abundance. (B) The mean atomic fraction of ^13^*C*, *f*_*b*_, for the dilution-corrected SIMS measurements (circles and diamonds) demonstrate inorganic carbon acquisition by the bacteria. Error bars correspond to the standard errors. (C) Bacterial growth measured using viable counts, plotted on a logarithmic scale as the mean (with standard error shown as error bars) of two measurements (circles and diamonds), shows an increase in the exponential growth rate and carrying capacity for a higher initial concentration of glycerol. The red crosses indicate the points that were unexpectedly high (approximately 1 × 10^12^ *cfu mL*^−1^) and therefore considered outliers and not included in the parameter optimisation. The results of the model fit, with parameters as specified in Supplementary Tables S6 and S7, are also plotted for the (B) atomic fraction *f*_*b*_ and (C) cell density *b*. For the 0.1 % glycerol culture the results from two different parameter optimisations are compared. For the fit that includes respiration (solid line), i.e. *η* included as a free parameter, the results are given in Supplementary Tables S6 and S7. For the fit that neglects bacterial respiration (dotted line), i.e. *η*′ = 1, the parameter optimisation results are *K*_*c*_ = 6.6 × 10^−6^ *molC mL*^−1^, *μ*_*b*_ = 0.15 *h*^−1^ and for the 0.1 % glycerol culture *b*(0) = 1.2 × 10^7^ *cells mL*^−1^ and *X* = 0.025.

Concomitantly to SIMS, bacterial abundance was quantified using viable counts. As expected, higher glycerol concentrations resulted in faster exponential growth and larger carrying capacities (Figure 2C). Even with no glycerol added to the growth medium, bacterial growth was still observed (see also Figure S7). This was likely due to internal stored carbon carried forward from the pre-culture. During the first 24 *h*, when all cultures analysed were in the exponential growth phase, greater ^13^*C*-enrichment was observed for bacteria grown with a higher concentration of glycerol (Figure 2B). Since only inorganic carbon was labelled, the increase in *f*_*b*_ demonstrates DIC acquisition by *M. loti*.

The co-culture model was applied to interpret the SIMS results for the axenic cultures of *M. loti*. Mathematically, the model used for axenic bacteria is given by equations (2), (3) and (4) in Materials and Methods, with *a* = *v* = *r*_*e*_ = *r*_*p*_ = 0, which describes logistic growth of a bacterial population growing on a limiting organic carbon source.

To fit the model to the SIMS and growth data, two global fits were performed, one including respiration and another ignoring it. In the latter case, the model was unable to reproduce the data well (dotted line in Figure 2B-C). This suggests that DIC uptake and respiration are essential to accurately describe the carbon kinetics of axenic bacteria. The bacteria grown with 0.1 % glycerol showed a prominent peak in *f*_*b*_, which the model without respiration was unable to reproduce (Figure 2B). This can be explained by considering that only respiration provides the feedback of unlabelled carbon necessary for *f*_*b*_ to decrease. Respiration converts glycerol to *CO*_2_, which is released into solution and lowers the total labelled fraction of DIC. Thus, the labelled fraction of carbon consumed by bacteria decreases, causing *f*_*b*_ to decrease. The fit results for the growth efficiency *η* ∈ [0.15 − 0.63] and DIC uptake parameter *X* ∈ [0.009 − 0.046] (Supplementary Table S7) are similar to those reported in the literature, e.g. *η* ∈ [0.05 − 0.6] (ref. 57) and *X* ∈ [0.014 − 0.065] (ref. 54). Moreover, the DIC uptake parameter *X* was found to increase as a function of the exponential growth rate *μ*_*B*_, according to *X* = *m* ln(*μ*_*B*_) + *n* with *m* = 0.0167 ± 0.0004, *n* = 0.0785 ± 0.0013 and *R*^2^ = 0.999. A negative correlation between the growth efficiency *η* and ln(*μ*_*B*_) was found, giving *η* = *p* ln(*μ*_*B*_) + *q* with *p* = −0.10 ± 0.12, *q* = 0.12 ± 0.36 and *R*^2^ = 0.282; see Supplementary Table S7 and Supplementary Figure S8. The increase in the DIC uptake parameter *X* can be reasonably associated with an increase in carboxylation reactions responsible for DIC acquisition with faster growth (54, 55), however a more detailed metabolic model would be required to further investigate the functional relationships emerging from our data.

Overall, this study of axenic cultures revealed how the combination of temporal SIMS measurements with modelling can help determine which key metabolic phenomena are responsible for observed isotope labelling dynamics.

### Carbon transfer from algae to bacteria in co-culture

To gain new insights into the establishment of mutualistic algal-bacterial interactions, we applied the combined SIMS-modelling approach to study a co-culture between *C. reinhardtii metE7* and *M. loti*. The algae were pre-labelled and not washed prior to co-culture inoculation (see Materials and Methods and Figure S9), therefore DOC from the pre-culture was carried over into the co-culture. This provided the best chance of observing bacterial assimilation of algal derived carbon, given that the time-scale for DOC to become available to bacteria in the co-culture had not been measured previously.

The labelled carbon kinetics in the co-culture were followed using SIMS over a period of 72 *h*. SIMS images (Figure 3A) were used to determine the atomic fraction of ^13^*C*, *f*, for individual bacterial and algal cells. As for axenic bacteria, the quantities *f*_*a*_ and *f*_*b*_ denote the average atomic fractions for a population of algae and bacteria respectively (Figure 3B); single cell heterogeneity is considered below. Sustained population growth was observed for both the algal and bacterial populations (Figure 3C), which implied that they were not nutrient limited. In spite of algal population growth, *f*_*a*_ remained approximately constant throughout the co-culture (Figure 3B), which indicates a likely equilibrium for ^13^*C* in algae, with *f*_*a*_ equal to the atomic fraction of DIC *f*_*i*_ (see equation (S30) in Supplementary Methods). The increase in *f*_*b*_ (Figure 3B) showed that the bacteria assimilated ^13^*C* compounds from the extracellular environment. However, on its own, the SIMS results could not provide information on the precise carbon kinetics within the co-culture. In the early stages of a co-culture the question remains: are cells growing on mutually produced nutrients, nutrients carried-over from pre-culture or internal stores? Combining SIMS data with our mechanistic model allowed this question to be addressed.

**Figure 3:**
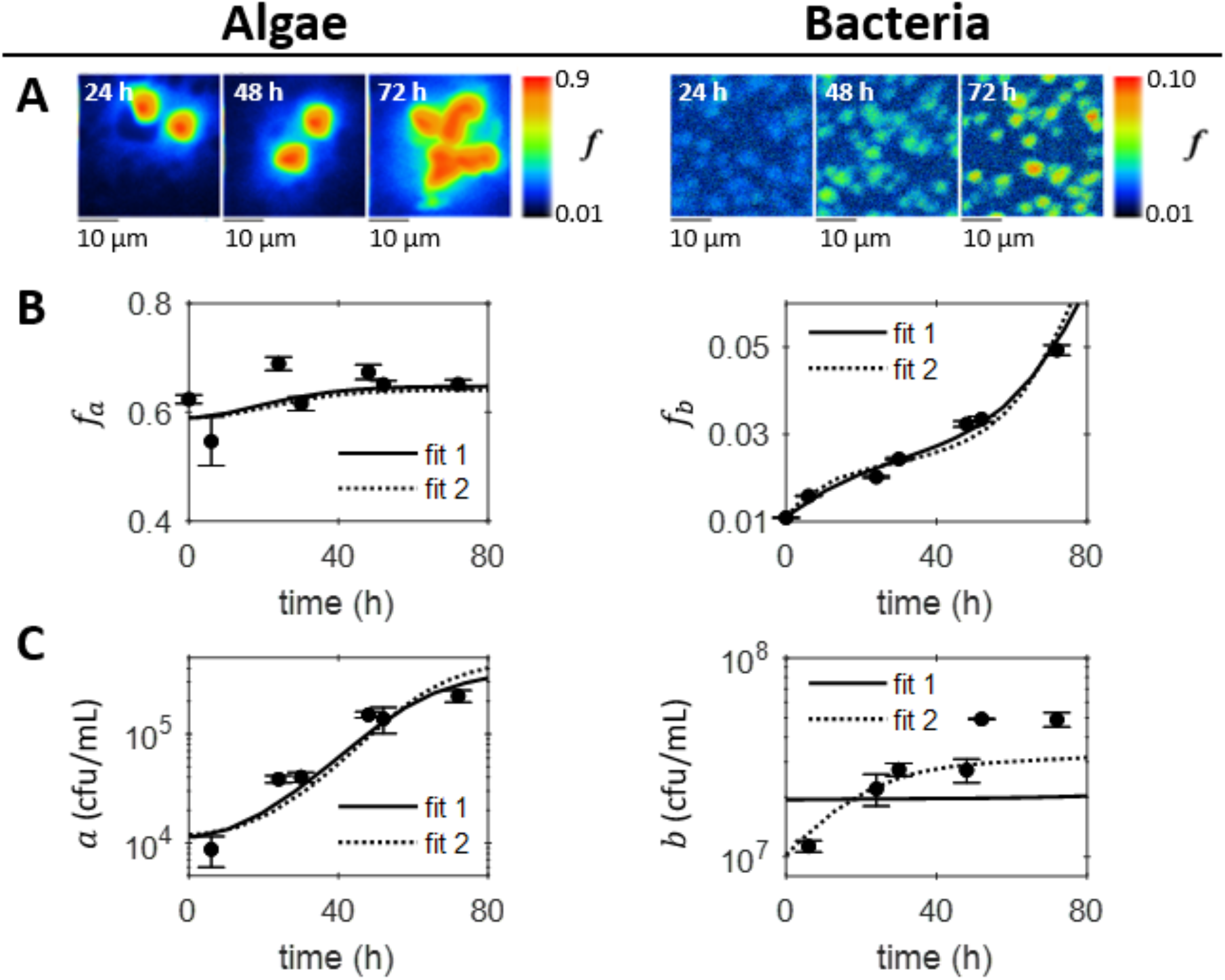
The algal-bacterial co-culture. (A) Example images of the atomic fraction of ^13^*C*, *f*, obtained by SIMS analysis of algal and bacterial cells sampled from the co-culture. The colour maps show the scale, starting at natural abundance. (B) The mean atomic fraction of ^13^*C*, *f*_*a*_ and *f*_*b*_ for algae and bacteria respectively, calculated from the dilution-corrected SIMS measurements for at least 5 algal cells and 100 bacterial cells per time-point (circles). Error bars correspond to the standard errors. (C) Algal and bacterial growth measured using viable counts, plotted as the mean (with standard error shown as error bars) of two viable count measurements (circles). The results of two model fits are also plotted for (B) the atomic fractions *f*_*a*_ and *f*_*b*_, and (C) cell densities *a* and *b*. Fit 1 fixed the initial *f*_*o*_(0) = 0.64, estimated using results for the pre-labelling culture of algae, whereas fit 2 included *f*_*o*_(0) as a free parameter. The model parameter values and initial conditions are as specified in Supplementary Tables S6 and S8. Although fit 2 gives a better fit to the data, it gives a low initial atomic fraction for the DOC *f*_*o*_(0) and high initial DOC concentration *c*_*o*_(0).

### Hidden nutrient kinetics

To further analyse the SIMS data and explore possible nutrient kinetics that couple the interaction partners, we formulated a mechanistic model of the algal-bacterial co-culture (see Materials and Methods) and performed parameter optimisations (see Supplementary Information, Figure S5, S6 and Tables S5 and S6). Figure 3B-C shows two separate global fits of the model to the algal and bacterial atomic fractions and cell densities. Fit 1 fixed the initial atomic fraction of ^13^*C* for DOC at *f*_*o*_(0) = 0.64, the expected value from the pre-labelled culture of algae (see Supplementary Information), whereas fit 2 included *f*_*o*_(0) as a free parameter. Fit 2 may appear to better describe the data, because it better reproduces bacterial growth, but the parameter optimisation result for *f*_*o*_(0) in fit 2 was close to natural abundance (Supplementary Table S8), which is not realistic for a culture expected to contain some labelled DOC from the highly labelled algal pre-culture. Neither fit was thus able to quantitatively capture the observations, suggesting that our model is probably too simple to be fully quantitative. Nonetheless, the model fits the data well qualitatively, and could be used to explore the nutrient kinetics that are not directly inferable from our measurements.

Using parameters from fit 1 (Supplementary Table S6), the model revealed the potential B_12_ and DOC kinetics driving the microbial growth dynamics (Figure 4A-B). The vitamin concentration *v* increases from zero (the co-culture medium was assumed to be initially vitamin-free because bacteria were washed thoroughly prior to establishing the co-culture and B_12_ was assumed to have been fully depleted in the pre-labelling culture of algae because it was inoculated with only 100 *ng L*^−1^ B_12_), and then starts to decrease after about 40 *h* (Figure 4A). Conversely, the DOC concentration *c*_*o*_ drops from the initial concentration *c*_*o*_(0), carried over from the unwashed algal pre-culture, and then starts to rise after approximately 30 *h* (Figure 4B), a few hours before the turnaround in B_12_ concentration. These results can be interpreted in terms of the production and consumption of nutrients, and the resulting population growth. At the start of the experiment bacterial DOC uptake during growth was likely responsible for the initial depletion of DOC (Figure 4B), which occurred at a faster rate than could be replenished by the algae. The model results also suggest that growing bacteria were initially producing B_12_ faster than the algal uptake rate, allowing the vitamin concentration to increase (Figure 4A). As it did so, the algae grew and photosynthesised, producing DOC to be utilised by the bacteria, which proliferated in turn. The turnaround in the nutrient kinetics occurs when production and consumption rates are matched, seen mathematically by setting 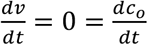 in equation (3). Figure 4A-B suggests that beyond the turning point at ≈ 30 *h*, DOC became more abundant as production by algae out-paced bacterial consumption. A short time later, the concentration of B_12_ began to decrease as production by bacteria fell below algal consumption.

**Figure 4:**
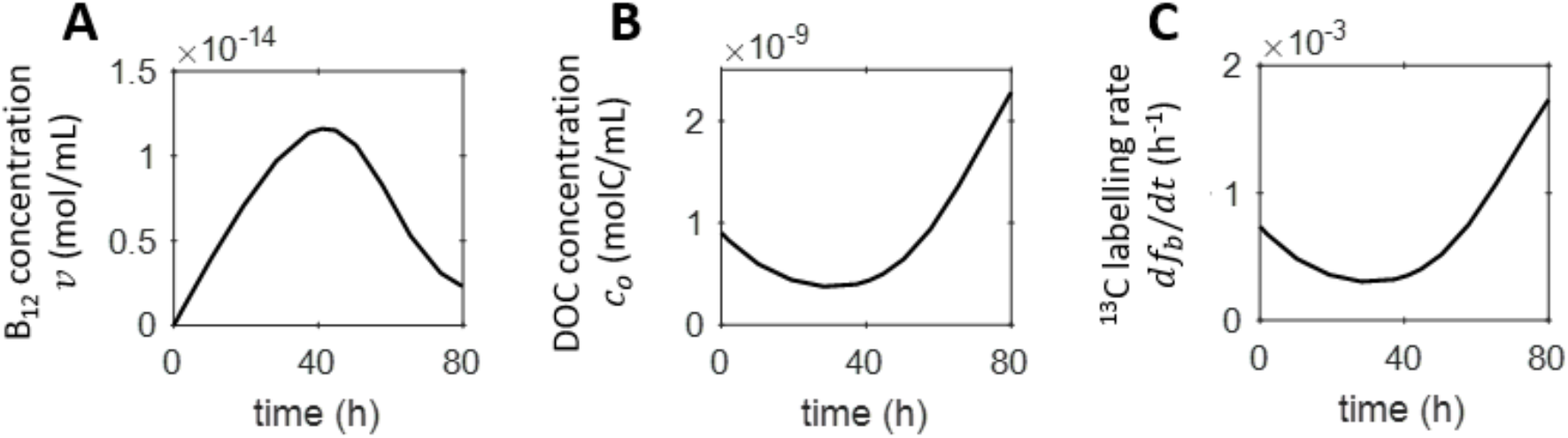
Nutrient kinetics in the co-culture predicted by the model. The concentrations of (A) B_12_ and (B) DOC in the co-culture predicted by the nutrient-explicit co-culture model using the parameter optimisation results obtained from fit 1, see Supplementary Table S6 for details of the parameter values and initial conditions used. (C) The isotope labelling rate calculated as 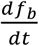 given by equation (4).

Furthermore, the time evolution of the derivative of the bacterial atomic fraction 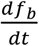 (obtainable from the model; equation (4)) is seen to mirror closely the fall and rise of the DOC, reproducing a turnaround at approximately the same time (Figure 4C). The model implies that this is because the rate of DOC uptake by bacteria is proportional to the DOC concentration, such that a decrease in the DOC concentration decreases the uptake rate, which directly slows down the rate of ^13^*C* assimilation. Thus, the model, while not providing a fully quantitative description of the growth dynamics, is nevertheless able to chart the temporal variation of the nutrient kinetics from isotope labelling experiments.

### Single cell heterogeneity

The SIMS results discussed thus far were averages obtained from several single cell measurements. We now turn to the heterogeneity in atomic fraction revealed by SIMS (see Supplementary Figures S3 and S4 for histograms of the single cell data). For this we concentrated on bacteria which provided better statistics than algae (minimum 80 bacterial cells measured per time point, versus 5 − 29 cells per time-point for algae). For unlabelled bacteria at natural abundance the single cell measurements showed a narrow distribution of atomic fractions (Supplementary Figure S3), indicating that all bacteria started at approximately the same value. For axenic bacteria, increasing the glycerol concentration caused greater DIC uptake, and ^13^*C* was seen to be more widely spread across the population (Supplementary Figure S3). For the highest glycerol concentration, the cell distribution was seen to broaden and then narrow again over time, corresponding to the rise and fall of the mean atomic fraction, and a transition of the culture to stationary phase. In contrast, for bacteria in co-culture, the distribution of single cell atomic fractions broadened steadily over time (Supplementary Figure S4).

These single cell results clearly indicate heterogeneity in isotope labelling across the bacterial populations. To analyse heterogeneity, a stochastic, structured model would strictly be required, for example as was used to explain how the circadian clock and environmental cycles affect cell size control and generate two subpopulations in the cyanobacterium *Synechococcus elongatus* (58). Our mean field model could, however, still be usefully applied to simulate heterogeneity and investigate potential origins of the observed single cell distributions by solving the model for parameter values above and below the fit results obtained for the mean atomic fractions (Supplementary Tables S6 and S7). Specifically, we considered the effect of varying the DIC uptake parameter *X*, bacterial maximum growth efficiency *η* and maximum bacterial growth rate *μ*_*b*_, with ranges given in the legend of Figure 5. The resulting variations in predicted bacterial atomic fractions (shaded areas in Figure 5) could then be compared with the variation observed experimentally, considered as the standard deviations of the SIMS single cell distributions (error bars in Figure 5). For axenic bacteria, a distribution in the values of *X* appeared to best account for the experimental standard deviation in the atomic fraction *f*_*b*_, especially for the culture grown at the highest glycerol concentration, where the model successfully reproduced the experimentally observed narrowing of the distribution at long times (Figure 5A). The comparison with experimental trends for variations in *η* and *μ*_*b*_ was less favourable (Figure 5B-C). Instead, for bacteria in co-culture, the progressive broadening of the distribution of *f*_*b*_ was best described by a distribution in *μ*_*b*_ (Figure 5C), with distributions in *η* and *X* not doing as well in the comparison (Figure 5A-B).

**Figure 5:**
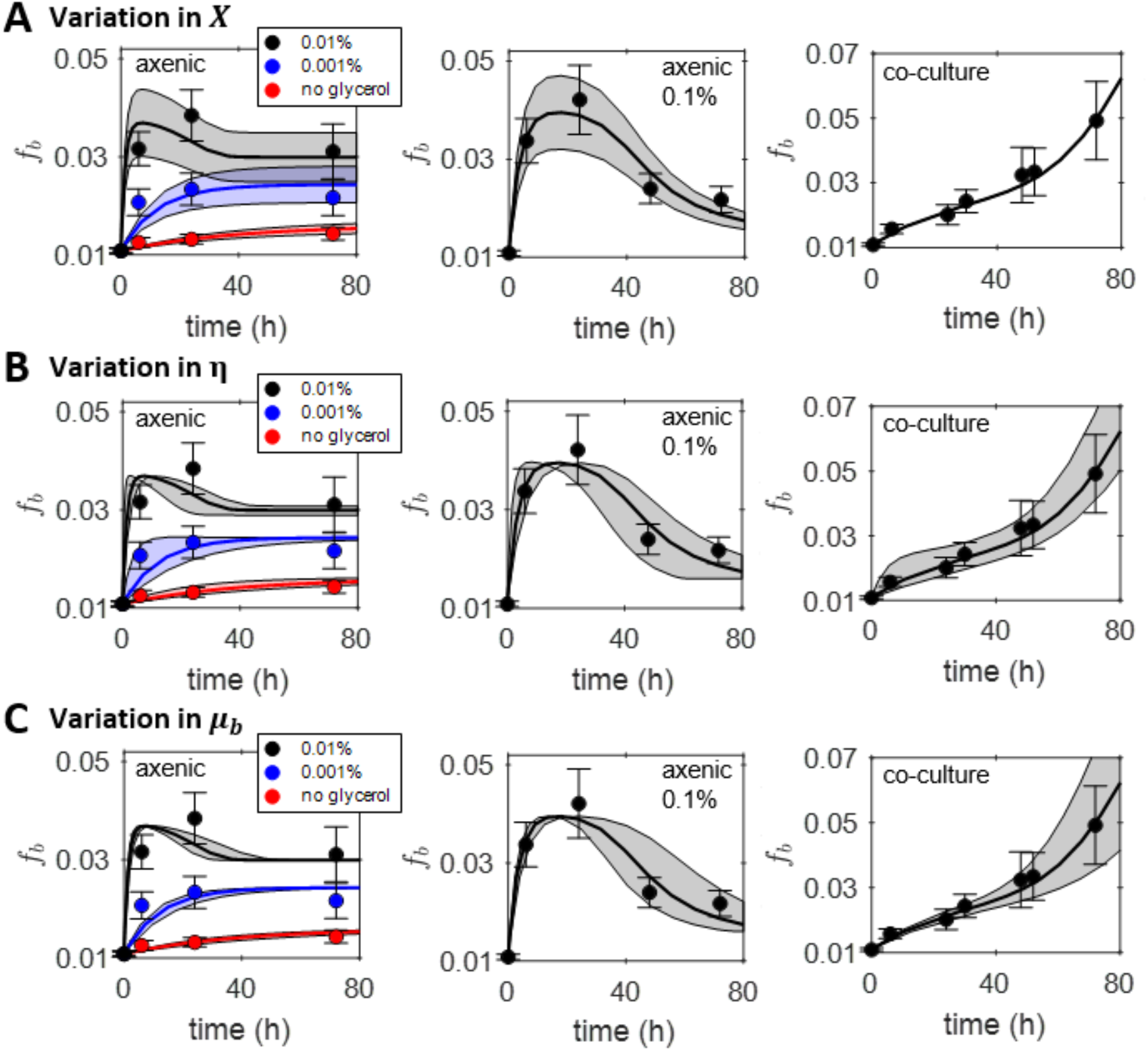
Comparison of single cell heterogeneity predicted by the model and measured experimentally with SIMS. The mean for the dilution-corrected results for *f*_*b*_ obtained using SIMS are plotted as circles with error bars indicating the standard deviation of the single cell values. The results of the model fit to the experiments is shown as a solid line and the shaded regions indicate the predicted range of *f*_*b*_ values when a range in a specific model parameter is considered. (A) The range of *X* values considered for the 0.1 %, 0.01 %, 0.001 % and no glycerol cultures of axenic bacteria were *X* ∈ [0.034,0.058], *X* ∈ [0.031,0.053], *X* ∈ [0.016,0.028] and *X* ∈ [0.007,0.011] respectively. For the co-culture the range considered was *X* ∈ [0.011,0.019]. (B) The range of *η* values considered for the 0.1 %, 0.01 %, 0.001 % and no glycerol cultures of axenic bacteria were *η* ∈ [0.21,0.81], *η* ∈ [0.05,0.25], *η* ∈ [0.09,0.69] and *η* ∈ [0.33,0.93] respectively. For the co-culture the range considered was *η* ∈ [0.11,0.91]. (C) For the axenic cultures of bacteria *μ*_*b*_ ∈ [0.11,0.19] and for the co-culture *μ*_*b*_ ∈ [0.34,0.50] in units *h*^−1^. Variation in *X* best accounts for the observed temporal trends in the standard deviations of the single cell data for the axenic cultures, whereas variation in *μ*_*b*_ best accounts for the co-culture results.

## Discussion

Whilst several studies have demonstrated mutualistic interaction between bacteria and algae mediated by nutrient exchange (43,44,46), none have integrated time-resolved SIMS with mechanistic modelling to elucidate nutrient kinetics, as we have done here. Our findings examine how nutrient kinetics control the inception and temporal development of an algal-bacterial mutualism. More broadly, this connects to the question of how co-occurrence can, on an evolutionary timescale, transform non-specialised relationships into more specialised partnerships, from streamlined microbial metabolisms (59–62) to plant-microbe interactions (63–65).

Initially, our SIMS-modelling approach demonstrated the uptake of labelled DIC by the heterotrophic bacterium *M. loti*, grown axenically on an unlabelled carbon source (glycerol). This confirmed similar results from previous studies of DIC uptake by heterotrophic bacteria (54, 55), while also providing more extensive data in terms of temporal dynamics and concentration of organic carbon. Fractional DIC uptake, described by the parameter *X*, and respiration, described by the bacterial growth efficiency parameter *η*, were essential for quantitatively describing the results. Fitting the model to results of ^13^*C* labelling experiments provided values for these parameters, an approach that could be used in future studies to investigate how these parameters are affected by environmental variables, including temperature, nutrient limitation and energetic quality of the organic carbon substrate (56, 66).

The SIMS-modelling approach was then used to shed light on the role of nutrient exchange during the onset of mutualistic interaction in a co-culture of *M. loti* bacteria and vitamin B_12_-dependent *C. reinhardtii metE7* algae. SIMS results showed that the bacteria assimilated algal-derived labelled carbon and using our mechanistic model we further revealed nutrient kinetics that couple the mutualistic partners. Initial DOC in the co-culture (carried forward from the algal pre-culture) delayed the onset of reciprocal mutualistic interaction: algae and bacteria started to grow exclusively on what each partner was producing only after about 30 *h* into the co-culture. A similar time-scale was observed in a NanoSIMS study of Antarctic microbial communities, which found that heterotrophic bacteria used organic carbon exudates from primary producers within 24 *h* (67).

Exploiting the single cell resolution of SIMS, our results revealed the heterogeneity of carbon uptake across a bacterial population. The distribution of atomic fractions for axenic bacteria displayed a width that was non-monotonic with time, whereas for the bacteria in co-culture with algae, this width increased monotonically. This difference in the temporal evolution of the standard deviation could be because DIC kinetics governed the isotope labelling in the axenic cultures, while the isotope labelling of co-cultured bacteria was likely dominated by uptake of algal derived DOC. To simulate variation of phenotypes across the bacterial population, our model was solved with parameter values above and below the fit results. A distribution in inorganic carbon uptake gave the best agreement with experiment for axenic cultures, whereas a distribution in bacterial growth rate best accounted for the co-culture measurements. This could well reflect the heterogeneous carbon environment for bacteria growing on algal exudates comprising a mix of compounds, each corresponding to a different growth rate. Conversely, axenic bacteria were fed on a single carbon substrate. Future studies could compare structured mechanistic models and computer simulations that describe variation in population dynamics and nutrient kinetics across microbial populations (58,68,69) with the approach to modelling heterogeneity used here.

Using a mechanistic model enhanced the interpretation of temporal nutrient kinetics data obtained using SIMS for an algal-bacterial co-culture. As discussed, the model we have constructed works well qualitatively, but comparison with the SIMS experiment points to possible improvements. For example, the model fit to SIMS data for the co-culture could benefit from better parametrisation of DOC production and its assimilation by bacteria. Further experiments that include DOC measurements would allow better estimates for the algal DOC export parameter *s*_*c*_ and the bacterial carbon uptake parameter *k*_*b*,*c*_ to be obtained. For the co-culture a discrepancy between bacterial growth and isotope labelling was observed, with estimates of net carbon assimilation rate from bacterial ^13^*C* enrichment measurements accounting for only about 6 % of bacterial population growth (see Supplementary Information). This suggests that the pre-cultured bacteria were not completely carbon starved and could grow using internal stores of organic carbon. Future models could account for internal carbon storage in bacteria, e.g. using nutrient kinetic models informed by flux balance analysis. Despite these limitations, the current model could be used to qualitatively predict mutualistic dynamics, e.g. how different species or mutant combinations would grow or how different initial conditions affect the interaction outcome. This could guide experimental investigation and accelerate discovery towards a mechanistic understanding of microbial interactions.

## Acknowledgements

H.L. acknowledges support from the EPSRC CDT in Nanoscience and Nanotechnology (NanoDTC), grant number EP/L015978/1. H.L. and O.A.C. gratefully acknowledge support from the Winton Programme for the Physics of Sustainability. O.A.C. and A.G.S. acknowledge EPSRC mobility fellowship EP/J004847/1. A.G.S. and F.B. acknowledge funding from BBSRC Doctoral Training Partnership BB/M011194/1. F.J.P. acknowledges support from Mines ParisTech and from a Raymond and Beverly Sackler Scholarship. We thank Howard Griffiths and Moritz Meyer for helpful discussions. We acknowledge James Rolfe and the Godwin Lab, Department of Earth Sciences, University of Cambridge for IRMS analysis. We acknowledge the KIVIK facility of Karolinska Institutet, and Andrea Caputo with assistance with the laser markings prior to SIMS. R.A.F. contribution, including the laser microscopy, was funded by an Academy Fellow Grant from the Knut and Alice Wallenberg Foundation. We thank Kerstin Lindén at the NordSIM facility at the Natural History Museum in Stockholm with assistance with sample preparation and SIMS analysis. The NordSIM facility acknowledges funding from the Swedish Museum of Natural History, Swedish Research Council (via infrastructure grant 2014-06375), University of Iceland and Consortium of Danish geoscience institutions.

## Author contributions

HL, OC, AGS and RF conceived the study. HL performed the experiments and the modelling, collected data, analysed data and solved the model. FJP and FB contributed experiments to parameterise the model. FJP assisted with the optimisation. HL, OC and AGS wrote the manuscript, which was commented on by all authors.

