## Supplementary Information for "Combining SIMS and mechanistic modelling to reveal nutrient kinetics in an algal-bacterial mutualism"

### I. Supplementary Figures and Tables

### II. Supplementary Methods

### III. Supplementary Results

### I. Supplementary Figures and Tables

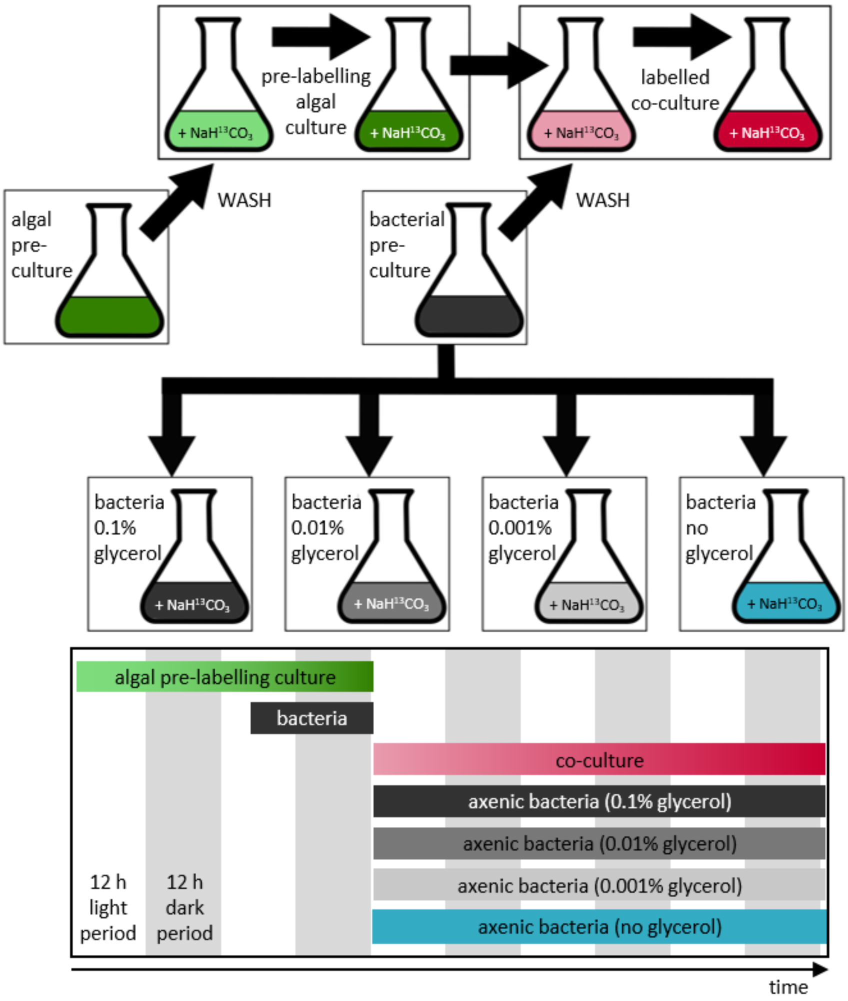

**Figure S1: Work-flow for the stable isotope labelling cultures.** Schematic overview and time-line of the stable-isotope labelling cultures using the alga *C. reinhartii metE7* and the bacterium *M. loti*, as described in detail in the text. The vertical white and grey bars indicate the 12 $h$ light and 12 $h$ dark periods respectively. Samples were taken at different time-points for single cell carbon isotope analysis using SIMS and bulk carbon isotope analysis of the algal and bacterial biomass using IRMS.

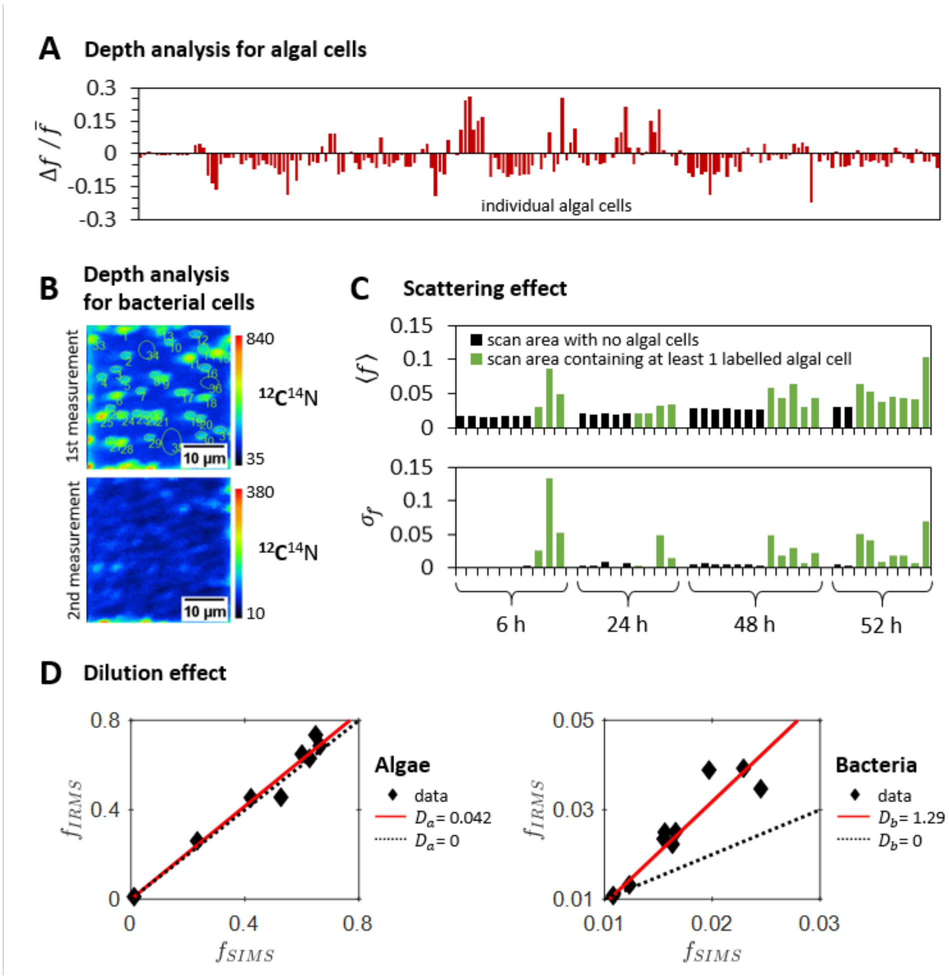

**Figure S2: Technical considerations.** (A) The difference between the atomic fraction of $^{13}C$ in algal cells obtained from the third and first measurements ($\Delta f = f_3 - f_1$) relative to the mean ($\bar{f} = (f_1 + f_2 + f_3)/3$). These results show that the carbon isotopes are not homogeneously distributed within the algal cells. (B) Example SIMS result for the $^{12}C^{14}N$ isotope images of bacteria obtained for two repeated measurements at the same sample location. The colour maps indicate the scale for the SIMS measurements in units of secondary ion counts, which were accumulated over 100 scans. These results imply that the majority of the bacterial biomass is sputtered away during the first measurement. (C) Comparison between the mean and standard deviation of the atomic fraction of $^{13}C$ in bacterial cells ($\langle f \rangle$ and $\sigma_f$ respectively) for scan areas of co-culture samples that do not contain any highly labelled algal cells (black bars) and areas that contain at least one highly labelled algal cell (green bars). These results imply a scattering effect causes the atomic fractions of $^{13}C$ obtained for bacterial cells to be both higher and more variable when the scan area contains a labelled algal cell.

(D) Atomic fraction of $^{13}C$ obtained by IRMS and SIMS analysis (black diamonds) for both algae (left) and bacteria (right), with the red lines showing the results of the least squares fit of equation (8) in Supplementary Methods, using $f_{ch} = 0.0108$ (Table S3). The $D = 0$ case is plotted (black dotted line) to show that if there was no dilution effect the IRMS and SIMS results would be expected to give the same results. The dilution effect means that the SIMS measurements provide an underestimate of the true, undiluted $f$.

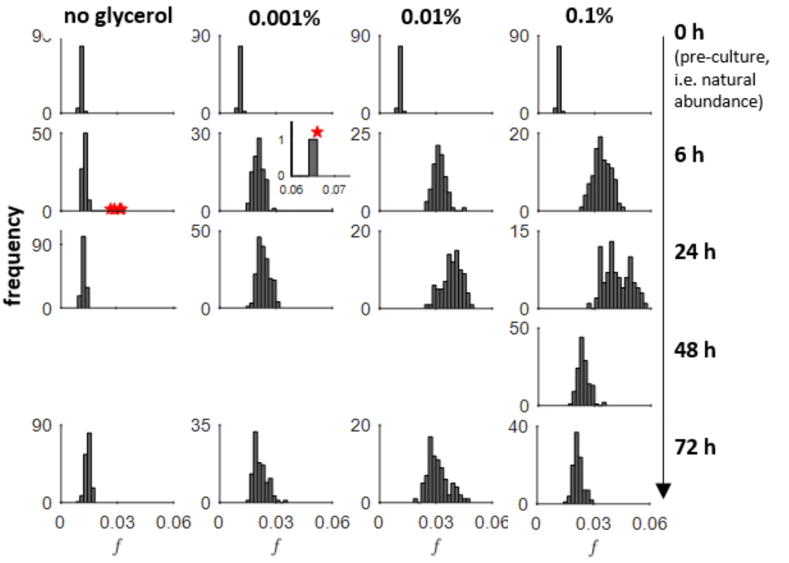

**Figure S3: Histograms for the SIMS results of bacterial cells grown in axenic cultures.** Histogram plots showing the dilution-corrected SIMS results for the single cell measurements of $f$. The red stars indicate the points that were considered outliers and therefore excluded from the calculation of the mean (i.e. $4$ points for the $6\ h$ sample from the no glycerol culture and $1$ point for the $6\ h$ sample from the $0.001\ \%$ glycerol culture, see Supplementary Methods for details).

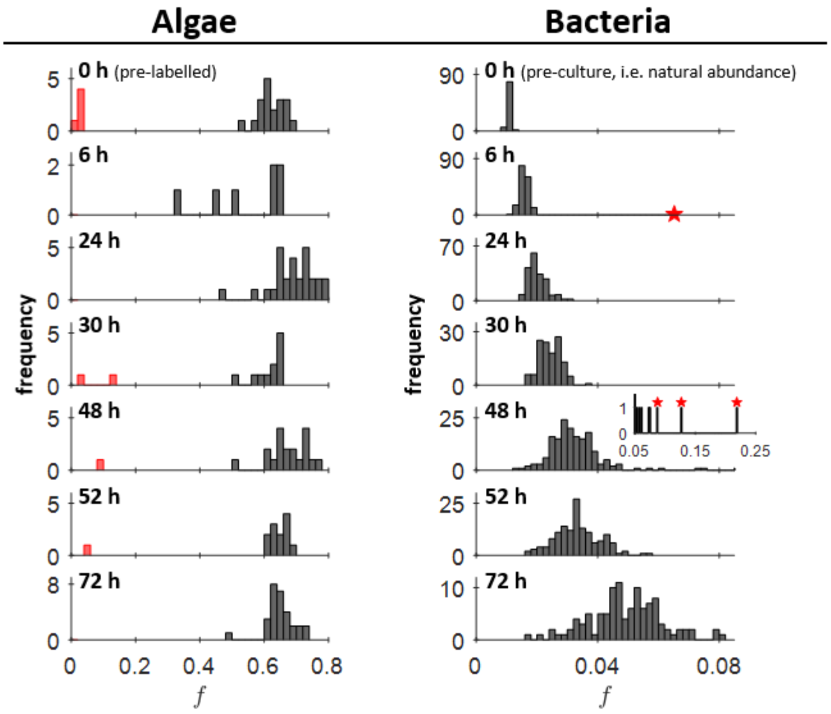

**Figure S4: Histograms for the SIMS results of algal and bacterial cells grown in co-culture.** Histogram plots showing the dilution-corrected SIMS results for single cell measurements of the atomic fraction of $^{13}C$ for algal and bacterial cells at different time-points of the co-culture. The red bars indicate the algal cells that were not included in the calculation of the mean because they were close to natural abundance and therefore considered inactive. The red stars indicate the bacterial cells that were considered outliers and therefore excluded from the calculation of the mean (see Supplementary Methods for details).

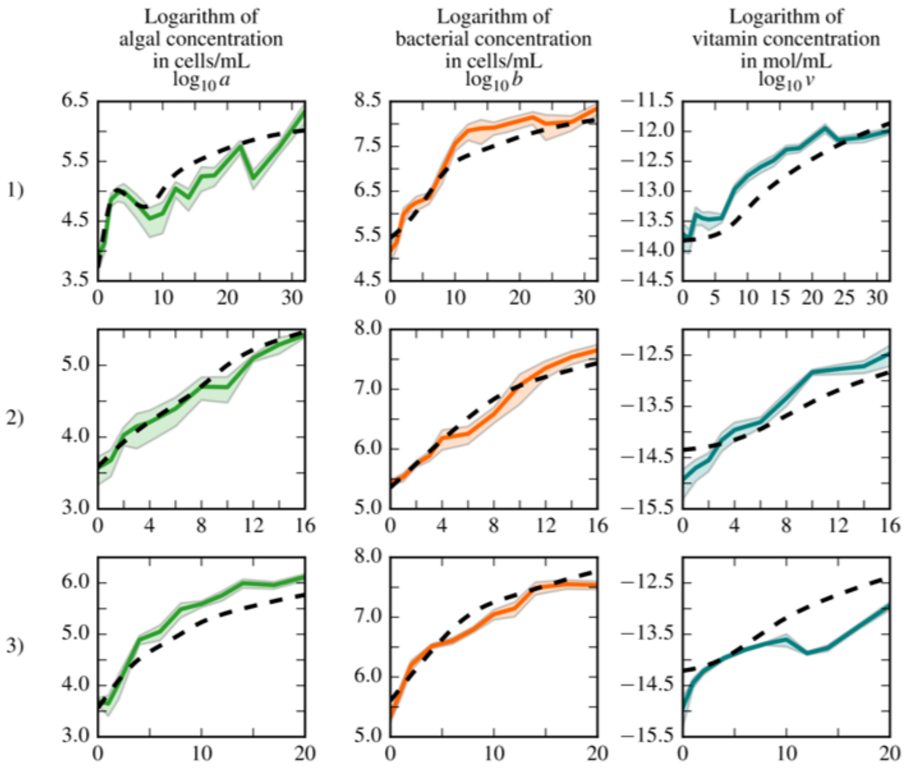

**Figure S5: Parameter optimisation for a simplified co-culture model.** Fit of data obtained for co-cultures of *C. reinhardtii metE7* and *M. loti*. Each row of plots corresponds to an independent experiment, with the first column the evolution of algal density, in the second column the evolution of bacterial density and in the third column the evolution of vitamin concentration as determined by bioassy. The mean of each variable appears as a continuous line, with the shaded region showing the standard deviation. The global fit with a unique set of parameters for the three independent experiments is shown in black dashed lines.

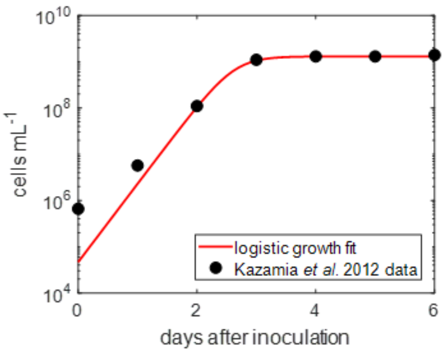

**Figure S6: Logistic growth fit for *M. loti*.** Data taken from Kazamia *et al.* (2012) (ref. 46) for *M. loti* grown axenically in 0.1 % glycerol was fit with the logistic growth equation $b = K_b/(1 + M\,e^{-r\,t})$, with $M = 2.8 \times 10^4$, $r = 3.9$ and carrying capacity $K_b = 1.3 \times 10^9$.

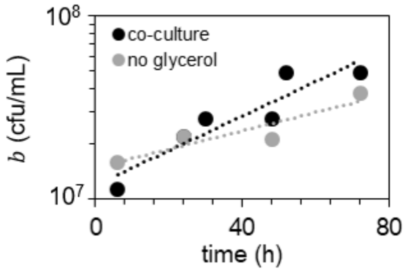

**Figure S7: Fit for the exponential growth rate of bacteria.** Plotted points show the viable count results for the growth of *M. loti* in the co-culture (black) and in the axenic culture grown without glycerol (grey). The dotted lines indicate the results for the exponential growth fit using equation $b = b(0)\exp(\mu_B\,t)$, giving $b(0) = 1.2 \pm 0.01 \times 10^7\ cfu\ mL^{-1}$ and $\mu_B = 0.022 \pm 0.005\ h^{-1}$ for the co-culture and $b(0) = 1.5 \pm 0.01 \times 10^7\ cfu\ mL^{-1}$ and $\mu_B = 0.012 \pm 0.004\ h^{-1}$ for the axenic culture.

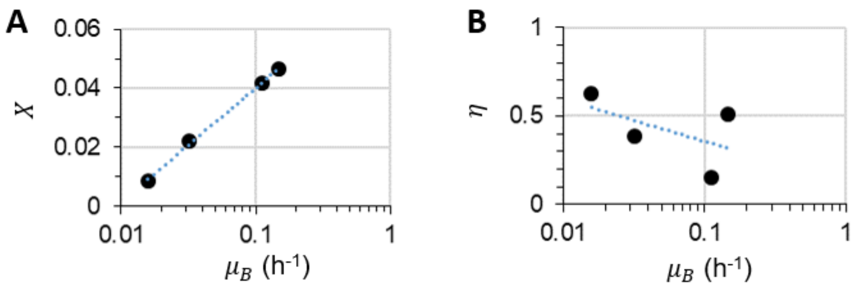

**Figure S8: For axenic bacteria the DIC uptake parameter and bacterial growth efficiency depend on the initial exponential growth rate.** (A) The relationship between the DIC uptake parameter $X$ and the initial exponential growth rate $\mu_B = \mu_b\,c_o(0)/(c_o(0) + K_c)$ was approximated with a logarithmic fit using equation $X = m\ln(\mu_B) + n$, giving $m = 0.0167 \pm 0.0004$ and $n = 0.0785 \pm 0.0013$, with $R^2 = 0.999$. (B) The relationship between the bacterial growth efficiency $\eta$ and $\mu_B$ was approximated with a logarithmic fit using equation $\eta = p\ln(\mu_B) + q$, giving $p = -0.10 \pm 0.12$ and $q = 0.12 \pm 0.36$, with $R^2 = 0.282$.

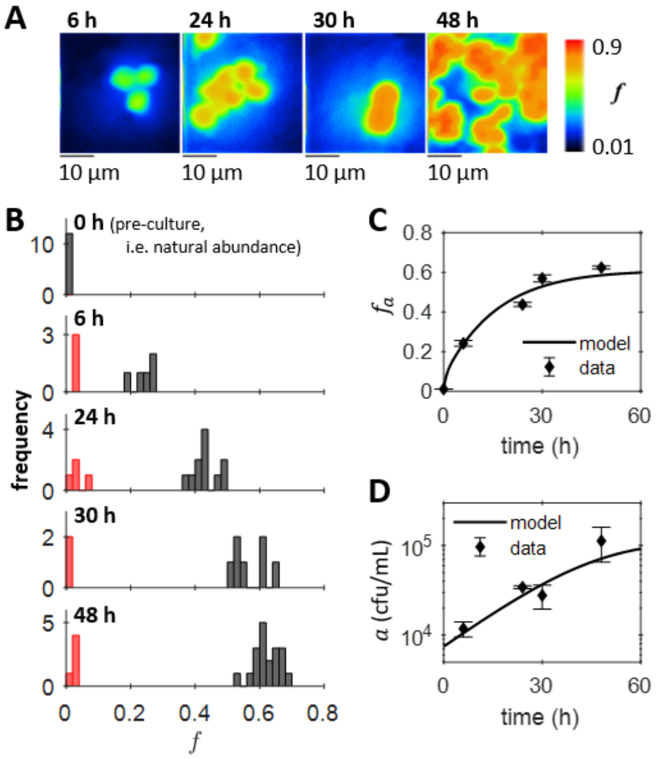

**Figure S9: The pre-labelling, axenic culture of *C. reinhardtii metE7*.** (A) Example images of the atomic fraction of $^{13}C$, $f$, obtained by SIMS analysis of algal cells sampled at different time-points of the pre-labelling, axenic culture grown with $5mM\ NaH^{13}CO_3$. The colour map shows the scale, starting at natural abundance. (B) Histogram plots showing the dilution-corrected SIMS results for single cell measurements of the atomic fraction of $^{13}C$ in individual algal cells. The red bars indicate the algal cells that were not included in the calculation of the mean because they were close to natural abundance and therefore considered inactive. (C) The mean atomic fraction of $^{13}C$ for the dilution-corrected SIMS measurements (diamonds). Error bars, showing the standard error, are small compared to the size of the plotted points. (D) Algal growth measured using viable counts ($cfu\ mL^{-1}$), plotted on a logarithmic scale as the mean and standard error of two measurements (diamonds). The results of the model fit, with parameters and initial conditions as specified in Table S6, are also plotted for (C) $f_a$ and (D) $a$.

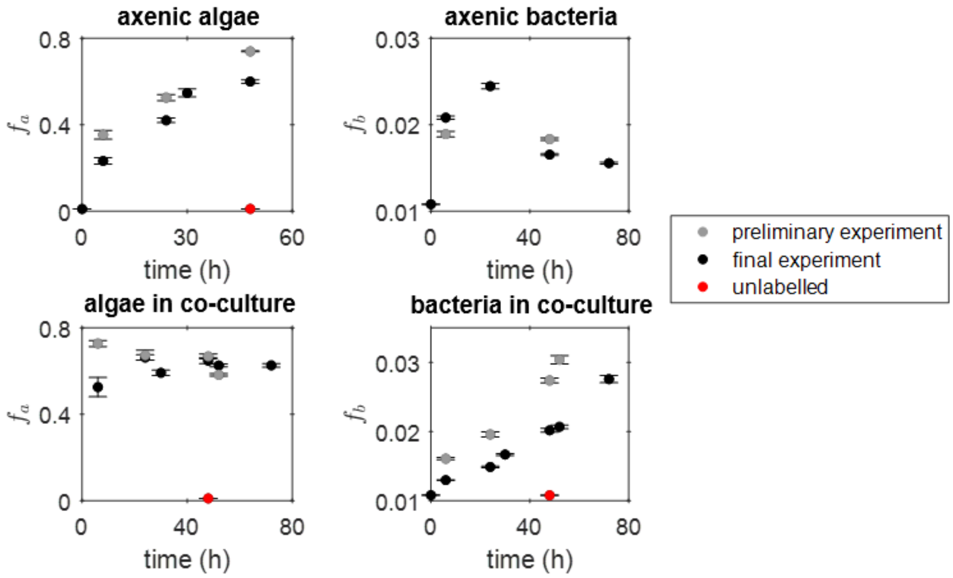

**Figure S10: Comparison of the SIMS results for the preliminary experiment, final experiment and unlabelled control cultures.** Preliminary SIMS results (grey circles) for the mean carbon isotope fraction are compared with the results from the final experiment (black circles) for the pre-labelling cultures of algae grown with 5 $mM\ NaH^{13}CO_3$, axenic cultures of bacteria grown with 0.1 % glycerol and 5 $mM\ NaH^{13}CO_3$ and the algal-bacterial co-culture. Error bars correspond to the standard errors. In the preliminary experiment, control cultures of axenic algae and a co-culture were grown with 5 $mM$ unlabelled $NaHCO_3$and a sample at 48 $h$ was taken and analysed using SIMS to show that the cells remained at natural abundance (red circles). For algal cells, the preliminary experiment only obtained one SIMS measurement, whereas for the final experiment the values plotted represent the mean value for algal cells where the single cell values are the mean of 2-3 repeated SIMS measurements taken at the same location on the filter. The SIMS results presented here have not been dilution-corrected.

**Supplementary Table S1: Trace elements adapted from Kropat *et al.* (2011) (ref. 50).** The concentrations of the different chemical components for each of the seven stock solutions of trace elements for the Tris-minimal media used in this work. For $1\ L$ of Tris-minimal media, $1\ mL$ of each solution was added.

| Number | Chemical Component | Concentration ($mM$) |
|---|---|---|
| 1 | $EDTA \cdot Na_2 \cdot 2H_2O$ | 25 |
| 2 | $(NH_4)_6Mo_7O_{24} \cdot 4H_2O$ | 0.032 |
| 3 | $CuCl_2 \cdot 2H_2O$<br>$EDTA$ | 1.4<br>2 |
| 4 | $ZnSO_4 \cdot 7H_2O$<br>$EDTA$ | 2.5<br>2.7 |
| 5 | $MnCl_2 \cdot 4H_2O$<br>$EDTA$ | 6<br>6 |
| 6 | $FeCl_3 \cdot 6H_2O$<br>$EDTA$<br>$Na_2CO_3$ | 20<br>22<br>22 |
| 7 | $CoCl_2 \cdot 6H_2O$ | 7 |

**Supplementary Table S2: List of cultures.** A complete list of the cultures grown as part of the stable isotope labelling experiments described in this work. Tris-minimal growth medium was used for all cultures with the addition of $B_{12}$, glycerol and sodium bicarbonate as listed in this table. These cultures were grown in $2\ L$ conical flasks except for the pre-cultures, which were grown in $1\ L$ flasks.

| Cultures for the preliminary experiment | Volume ($mL$) | $B_{12}$ ($ng/L$) | Glycerol ($\%v/v$) | Sodium bicarbonate |
|---|---|---|---|---|

| | | | | |
|---|---|---|---|---|
| Algal pre-culture | 600 | 1000 | | |
| Axenic algae (pre-labelling) | 1000 | 100 | | $5\ mM\ NaH^{13}CO_3$ |
| Axenic algae (unlabelled) | 1000 | 100 | | $5\ mM\ NaHCO_3$ |
| Bacterial pre-culture | 400 | | 0.1 | |
| Axenic bacteria (0.1% glycerol) | 1000 | | 0.1 | $5\ mM\ NaH^{13}CO_3$ |
| Labelled co-culture | 1000 | | | $5\ mM\ NaH^{13}CO_3$ |
| Unlabelled co-culture | 1000 | | | $5\ mM\ NaHCO_3$ |
| **Cultures for the final experiment** | **Volume ($mL$)** | **$B_{12}$ ($ng/L$)** | **Glycerol ($\%v/v$)** | **Sodium bicarbonate** |
| Algal pre-culture | 600 | 1000 | | |
| Axenic algae (pre-labelling) | 1000 | 100 | | $5\ mM\ NaH^{13}CO_3$ |
| Bacterial pre-culture | 400 | | 0.1 | |
| Axenic bacteria (0.1% glycerol) | 1000 | | 0.1 | $5\ mM\ NaH^{13}CO_3$ |
| Axenic bacteria (0.01% glycerol) | 1000 | | 0.01 | $5\ mM\ NaH^{13}CO_3$ |
| Axenic bacteria (0.001% glycerol) | 1000 | | 0.001 | $5\ mM\ NaH^{13}CO_3$ |
| Axenic bacteria (no glycerol) | 1000 | | | $5\ mM\ NaH^{13}CO_3$ |
| Labelled co-culture | 1000 | | | $5\ mM\ NaH^{13}CO_3$ |

**Supplementary Table S3: The dilution factor results.** The dilution factor, $D$, was obtained from a least squares fit of equation (8) in Supplementary Methods using the curve fitting application in Matlab and with $f_{ch} = 0.0108$. This table lists the results for $D$, the 95 % confidence bounds, the number of points in the fit, $n$, and the least square displacements, $R^2$. For bacteria, the fit was carried out using only data from axenic cultures.

| | $D$ | 95% confidence bound | $n$ | $R^2$ |
|---|---|---|---|---|
| **Algae** | 0.04 | $\pm 0.07$ | 8 | 0.968 |
| **Bacteria** | 1.29 | $\pm 0.41$ | 9 | 0.84 |

**Supplementary Table S4: C-N content and carbon yield.** Summary of the IRMS results for the carbon and nitrogen content and the carbon yield for the alga *C. reinhardtii metE7* and bacterium *M. loti*. The table gives the mean, standard error and the number of samples included in the mean ($n$).

| | | Mean | Standard error | $n$ |
|---|---|---|---|---|
| **Algae** | $\%AmtC$ | 35 | 4 | 8 |
| | $\%AmtN$ | 8 | 1 | 8 |
| | $C:N$ ratio | 4.4 | 0.1 | 8 |
| | $Y_{a,c}$ $(cells\ molC^{-1})$ | $4 \times 10^{12}$ | $1 \times 10^{12}$ | 4 |
| **Bacteria** | $\%AmtC$ | 39 | 2 | 13 |
| | $\%AmtN$ | 11 | 1 | 13 |
| | $C:N$ ratio | 3.71 | 0.03 | 13 |
| | $Y_{a,c}$ $(cfu\ molC^{-1})$ | $5 \times 10^{14}$ | $1 \times 10^{14}$ | 10 |

**Supplementary Table S5: Boundary conditions for the parameter optimisations.** These were the boundary conditions used for the various free parameters and free initial conditions of the parameter optimisations run for axenic algae, axenic bacteria and the co-culture. When units are not specified the parameter/initial condition is in dimensionless units. The boundary conditions for $\phi_S$, $\eta$ and $X$ came from their definition requiring these parameters to be between 0 and 1. Other boundary conditions were chosen to ensure that the parameter optimisation results were reasonable when considering their biological interpretation.

| | **Boundary Condition** | **Units** |
|---|---|---|
| All parameter optimisations: | $0 \leq \varphi_s \leq 0.99$<br>$0.01 \leq \eta \leq 1$<br>$0 \leq X \leq 1$ | |
| Axenic algae: | $0 \leq s_c \leq 10$<br>$0 \leq \hat{v}(0) \leq 5$<br>$0.001 \leq \hat{a}(0) \leq 0.01$<br>$0 \leq f_i(0) \leq 1$ | |
| Axenic bacteria: | $5 \times 10^6 \leq b(0) \leq 5 \times 10^7$<br>$0.01 \leq \mu_b \leq 2$<br>$1 \times 10^{-10} \leq K_c \leq 1 \times 10^{-4}$ | $cells\ mL^{-1}$<br>$h^{-1}$<br>$molC\ mL^{-1}$ |
| Axenic bacteria, no glycerol: | $0 \leq c_o(0) \leq 4 \times 10^{-7}$ | $molC\ mL^{-1}$ |
| Co-culture: | $0 \leq s_c \leq 10$<br>$0.001 \leq \hat{a}(0) \leq 0.01$<br>$0.001 \leq \hat{b}(0) \leq 0.03$<br>$1 \times 10^{-5} \leq \hat{c}_o(0) \leq 0.5$<br>$0.0108 \leq f_o(0) \leq 1$ | |

**Supplementary Table S6: Model parameters and initial conditions.** Full set of model parameters and initial conditions for *C. reinhardtii metE7* and *M. loti* grown both axenically and in co-culture.

| **Parameter** | **Symbol** | **Units** | **Axenic algae** | **Axenic bacteria** | **Co-culture** |
|---|---|---|---|---|---|
| Algal carrying capacity | $K_a$ | $cells\ mL^{-1}$ | $2.3 \times 10^6$ [a] | | $2.3 \times 10^6$ [a] |
| Bacterial carrying capacity | $K_b$ | $cells\ mL^{-1}$ | | $1.3 \times 10^9$ [b] | $1.14 \times 10^9$ [a] |
| B12 half-saturation concentration | $K_v$ | $mol\ mL^{-1}$ | $2.6 \times 10^{-14}$ [a] | | $2.6 \times 10^{-14}$ [a] |
| DOC half-saturation concentration | $K_c$ | $mol\ mL^{-1}$ | | $1.5 \times 10^{-6}$ [c] | $6.3 \times 10^{-7}$ [d] |
| Maximum bacterial growth rate | $\mu_b$ | $h^{-1}$ | | 0.15 [c] | 0.42 [a] |
| Maximum algal growth rate | $\mu_a$ | $h^{-1}$ | 0.21 [a] | | 0.21 [a] |
| Algal B12 yield | $Y_{a,v}$ | $cells\ mol^{-1}$ | $1.13 \times 10^{19}$ [e] | | $1.13 \times 10^{19}$ [e] |
| Algal carbon yield | $Y_{a,c}$ | $cells\ mol^{-1}$ | $4 \times 10^{12}$ [f] | | $4 \times 10^{12}$ [f] |
| Bacterial carbon yield | $Y_{b,c}$ | $cells\ mol^{-1}$ | | $5 \times 10^{14}$ [f] | $5 \times 10^{14}$ [f] |
| Per cell B12 production rate | $p_v$ | $mol\ cell^{-1}\ h^{-1}$ | | | $2 \times 10^{-23}$ [g] |
| DOC production rate | $p_c$ | $mol\ cell^{-1}\ h^{-1}$ | | $5.5 \times 10^{-15}$ [h] | |
| Fraction of storage | $\phi_s$ | | 0.87 [i] | | 0.9 [j] |
| Maximum BGE | $\eta$ | | | * | 0.51 [j] |
| DIC uptake fraction | $X$ | | | * | 0.015 [j] |
| **Non-dimensional parameter** | **Symbol** | **Definition** | **Axenic algae** | **Axenic bacteria** | **Co-culture** |
| Ratio of maximum growth rates | $\varepsilon$ | $= \mu_a/\mu_b$ | 0.51 [a] | | 0.51 [a] |
| Algal B12 uptake parameter | $k_{a,v}$ | $= K_a/(K_v\ Y_{a,v})$ | 7.8 [a] | | 7.8 [a] |
| Algal carbon uptake parameter | $k_{a,c}$ | $= K_a/(K_c\ Y_{a,c})$ | 0.91 [k] | | 0.91 [k] |
| Bacterial carbon uptake parameter | $k_{b,c}$ | $= K_b/(K_c\ Y_{b,c})$ | | 1.73 [l] | 3.6 [a] |
| B12 production strength | $s_v$ | $= (p_v\ K_b)/(\mu_a\ K_v)$ | | 4.2 [a] | 4.2 [a] |

| | | | | | |
|---|---|---|---|---|---|
| DOC production strength | $s_c$ | $= (p_c\, K_a)/(\mu_b\, K_c)$ | | 2.13 [i] | 0.047 [m] |

| **Initial conditions** | **Symbol** | | **Axenic algae** | **Axenic bacteria** | **Co-culture** |
|---|---|---|---|---|---|
| Algal cell density | $\hat{a}(0)$ | | 0.0032 [i] | 0 | 0.005 [m] |
| Bacterial cell density | $\hat{b}(0)$ | | 0 | * | 0.017 [m] |
| DOC concentration | $\hat{c}_o(0)$ | | 0 | * | 0.0014 [m] |
| B12 concentration | $\hat{v}(0)$ | | 0.374 [i] | 0 | 0 |
| DIC concentration | $\hat{c}_i(0)$ | | 5 | 5 | 5 |
| Algal atomic fraction | $f_a(0)$ | | 0.0108 | | 0.59 [n] |
| *Photosynthetically-active* atomic fraction | $f_{a,p}(0)$ | | 0.0108 | | 0.65 [n] |
| Bacterial atomic fraction | $f_b(0)$ | | 0.0108 | 0.0108 | 0.0108 |
| DOC atomic fraction | $f_o(0)$ | | 0.0108 | 0.0108 | 0.64 [n] |
| DIC atomic fraction | $f_i(0)$ | | 0.65 [i] | 0.65 [o] | 0.65 [o] |

* See Supplementary Table S7.

[a] From fitting a simplified co-culture model (i.e. $\phi_s = 0, \eta' = 1$ and $X = 0$) to population growth and $B_{12}$ concentration data, see Supplementary Methods for details.

[b] From fitting a logistic growth equation to data obtained by Kazamia *et al.* (2012) (ref. 46) for *M. loti* grown axenically with 0.1 % glycerol, see Supplementary Methods for details.

[c] From a global parameter optimisation performed for the four axenic cultures of *M. loti* grown with different concentrations of glycerol, see Supplementary Methods for details. The residual sum of squares for this global parameter optimisation result was 0.58, whereas when respiration was not included in the model it was 2.24.

[d] From the definition $K_c = \frac{K_b}{Y_{b,c}\, k_{b,c}}$.

[e] From the definition $Y_{a,v} = \frac{K_a}{K_v\, k_{a,v}}$.

[f] From dry mass measurements and IRMS analysis, see Supplementary Materials for details.

[g] From the definition $p_v = \frac{s_v \, \mu_a \, K_v}{K_b}$.

[h] From the definition $p_c = \frac{s_c \, \mu_b \, K_c}{K_a}$.

[i] Parameter optimisation results from fitting the model to the axenic, pre-labelling culture of *C. reinhardtii metE7*, see Supplementary Methods for details. The residual sum of squares for this global parameter optimisation result was 0.313. For comparison, when storage was not included in the model (i.e. $\phi_s = 0$), the parameter optimisation result gave $s_c = 0.041$ and the residual sum of squares was 0.323.

[j] Estimates obtained using parameter optimisation results for the axenic cultures, see Supplementary Methods for details.

[k] From the definition $k_{a,c} = \frac{K_a}{K_c \, Y_{a,c}}$.

[l] From the definition $k_{b,c} = \frac{K_b}{K_c \, Y_{b,c}}$.

[m] Parameter optimisation results from fitting the model to co-culture growth and SIMS data, i.e. fit 1 in Supplementary Table S8, see Supplementary Methods for further details.

[n] Estimates obtained using the model results for the axenic, pre-labelling culture of algae, see Supplementary Results.

[o] From the parameter optimisation result for the axenic, pre-labelling culture of algae, see Supplementary Results.

**Supplementary Table S7: Culture specific model parameters and initial conditions for axenic bacteria.** Model parameter values for the axenic cultures of *M. loti* grown with different concentrations of glycerol determined by a global parameter optimisation performed for the four axenic cultures of *M. loti* grown with $0.1$ %, $0.01$ %, $0.001$ % and no glycerol. The global free parameters were $\mu_b$ and $K_c$, which were constrained to have the same value for all four cultures. The free parameters and initial conditions that were permitted to be different for the different cultures were $\eta$, $X$ and $b(0)$. The initial DOC concentration $c_o(0)$ for the culture grown without glycerol was also included as a free parameter. The fixed initial conditions were $\hat{c}_i(0) = 5$, $\hat{v}(0) = 0$, $f_b(0) = 0.0108$, $f_o(0) = 0.0108$ and $f_i(0) = 0.65$, since for the experiments it was assumed that the DIC was in excess, initially there was no $B_{12}$ and the bacteria had natural abundance, the glycerol was unlabelled and the atomic fraction of $^{13}C$ in the DIC was taken as the estimate obtained from the parameter optimisation for axenic algae (see Supplementary Table S6). The residual sum of squares for this global parameter optimisation result was 0.58, whereas when respiration was not included in the model it was 2.24.

| Culture | $\boldsymbol{c_o(0)}$ $\boldsymbol{(molC\ mL^{-1})}$ | $\boldsymbol{b(0)}$ $\boldsymbol{(cells\ mL^{-1})}$ | $\boldsymbol{\eta}$ | $\boldsymbol{X}$ |
|---|---|---|---|---|
| 0.1 % glycerol | $4 \times 10^{-5}$ [a] | $8.8 \times 10^{6}$ | 0.51 | 0.046 |
| 0.01 % glycerol | $4 \times 10^{-6}$ [a] | $1.6 \times 10^{7}$ | 0.15 | 0.042 |
| 0.001 % glycerol | $4 \times 10^{-7}$ [a] | $1.8 \times 10^{7}$ | 0.39 | 0.022 |
| no glycerol | $1.7 \times 10^{-7}$ | $1.3 \times 10^{7}$ | 0.63 | 0.009 |

[a] Not free in the parameter optimisation, calculated from the % glycerol concentration using the molar mass of glycerol, 92.09 $g\ mol^{-1}$, and its density, 1.26 $g\ mol^{-1}$.

**Supplementary Table S8: Comparison of parameter optimisation results for the algal-bacterial co-culture.** Results of different parameter optimisation results for the co-culture between *C. reinhardtii metE7* and *M. loti*. The only free parameter was $s_c$ and the free initial conditions were $\hat{a}(0)$, $\hat{b}(0)$ and $\hat{c}_o(0)$, with $\hat{f}_o(0)$ included as an additional free initial condition for fit 2. The parameters $\phi_s = 0.9$, $\eta = 0.51$ and $X = 0.015$ were estimated using results from axenic cultures as specified in the text. All other parameters had values as specified in table S6. The fixed initial conditions were $\hat{c}_i(0) = 5$ (i.e. DIC concentration in excess), $\hat{v}(0) = 0$ (i.e. initially no $B_{12}$ in the media), $f_a(0) = 0.59$ and $f_{a,p}(0) = 0.65$ (i.e. using model results for the pre-labelling, axenic culture of algae, see text for details), $f_b(0) = 0.0108$ (i.e. bacteria initially have natural abundance), and $f_i(0) = 0.65$ (i.e. from the parameter optimisation result for axenic algae, see Supplementary Table S6).

| **Fit** | $s_c$ **DOC export parameter** | $\hat{a}(0)$ **Initial algal cell density** | $\hat{b}(0)$ **Initial bacterial cell density** | $\hat{c}_o(0)$ **Initial DOC concentration** | $f_o(0)$ **Initial DOC atomic fraction** | $r^2$ **Residual sum of squares** |
|---|---|---|---|---|---|---|
| 1 [a] | 0.047 | 0.005 | 0.017 | 0.0014 | 0.64 [a] | 1.96 |
| 2 [b] | 0.074 | 0.005 | 0.009 | 0.13 | 0.0144 | 1.29 |

[a] Initial atomic fraction of $^{13}C$ for the DOC, $f_o(0) = 0.64$, estimate obtained using the parameter optimisation result for axenic algae.

[b] Initial atomic fraction of $^{13}C$ for the DOC included as a free parameter.

### II. Supplementary Methods

#### IRMS analysis and estimating the algal and bacterial carbon yield

##### *Sample preparation and analysis*

To prepare the IRMS samples, first the biomass was concentrated using centrifugation. For co-cultures, an additional slow centrifugation step was used to concentrate the algal cells as a pellet, the supernatant was then passed through a 3 $\mu m$ filter and the filtrate was used as the bacterial fraction of the co-culture biomass, which was concentrated into a pellet by centrifugation. The concentrated biomass samples were transferred to eppendorfs and dried overnight in an oven at $50°C$. To remove any excess $NaH^{13}CO_3$ the samples were placed in a desiccator with 32 % $HCl$ for acid fumigation.

The dry mass of the samples was measured and the required amount for IRMS analysis was weighed out and encapsulated in tin. It was not possible to collect enough dry mass for IRMS analysis at every time-point. For samples with enough dry mass, $1$ to $4$ sub-samples were analysed at the Godwin lab, Department of Earth Sciences, University of Cambridge using the Thermo Delta V Plus and Costech.

#### *Estimating the algal and bacterial carbon yield*

The carbon and nitrogen content of *M. loti* and *C. reinhardtii metE7* were obtained from IRMS analysis, the results are given in Table S4. To calculate the carbon yield for algal cells (i.e. $Y_{a,c}$, the number of cells per mole of carbon) we used the equation

$$Y_{a,c} = M_r \frac{a \cdot V \cdot 100}{m \cdot \%\mathrm{AmtC}}, \quad (1)$$

with $a$ the algal cell density in $cells\ mL^{-1}$ measured using a Coulter counter, $V$ the sample volume in $mL$, $m$ the sample dry mass in $g$, $\%\mathrm{AmtC}$ the percent of dry mass that is carbon and $M_r$ the molar mass of carbon in $g\ mol^{-1}$. The value for the molar mass depends on the atomic fraction of $^{13}C$ and therefore $M_r$ was calculated using

$$M_r = 12 \cdot (1 - f) + 13 \cdot f, \quad (2)$$

with $f$ the atomic fraction of $^{13}C$ obtained from IRMS analysis. There were four algal samples that had suitable dry mass and IRMS measurements to be able to estimate the carbon yield. From these four estimates the carbon yield for algal cells was found to be $4 \pm 1 \times 10^{12}\ cells\ molC^{-1}$. The carbon yield for bacterial cells, $Y_{b,c}$, was calculated in the same way, but using the viable count measurement of cell density in $cfu\ mL^{-1}$. There were ten bacterial samples that had suitable dry mass and IRMS measurements to be able to estimate the carbon yield. From these ten estimates the carbon yield for bacterial cells was found to be $5 \pm 1 \times 10^{14}\ cells\ molC^{-1}$.

### SIMS technical details

#### *Sample preparation*

***Chemical fixation with formaldehyde***. For every $10\,mL$ of sample volume, $0.54\,mL$ of $37-41\,\%\,(w/v)$ formaldehyde was added to reach a final formaldehyde concentration of about $2\,\%\,(w/v)$. The sample was gently vortexed and then incubated at $4-6°C$ for $1\,h$. To remove the fixative, the sample was washed twice by centrifugation followed by re-suspension in 1X PBS buffer (i.e. phosphate buffered saline solution consisting of $10\,mM\ Na_2HPO_4$ and $150\,mM\ NaCl$). The sample was then centrifuged for a third time and finally re-suspended in a $1:1$ by volume mix of 1X PBS buffer and 96 % ethanol solution. Samples were stored in the fridge ($4-6°C$) until further use.

***Cell staining and vacuum filtration.*** In order to be able to visualise the distribution of algal and bacterial cells on the membrane filter, SYTO9 green fluorescent nucleic acid stain (taken from a Molecular Probes LIVE/DEAD BacLight bacterial viability kit) was used for both bacterial and algal cells. Per $1\,mL$ of sample, $1.5\,\mu L$ of $3.34\,mM$ SYTO9 was added, the sample was then incubated in the dark and at room temperature for 15 minutes. An appropriate sample volume was chosen for vacuum filtration in order to achieve an even distribution of cells on the filter, which meant choosing a volume that contained $0.5\times10^5$ to $2\times10^5\,cfu$ for algae and $1\times10^7$ to $1\times10^8\,cfu$ for bacteria. Isopore membrane filters with a pore size of $0.22\,\mu m$ and diameter $25\,mm$ (Merck Millipore) were pre-sputtered with $\approx 20\,nm$ gold coating, using a BioRad SEM Coating System, and cells were then deposited on these gold-coated filters by vacuum filtration using a Charles Austen Capex 8C vacuum pump.

***Confocal microscopy.*** It is important that samples prepared for SIMS are flat, as an uneven sample can result in unreliable measurements (1). An Olympus Fluoview laser scanning confocal microscope (FV1200) was used to image the filter samples and to ensure an even distribution of cells. A $473\,nm$

excitation laser was used and fluorescence emission was detected in two channels; $490 - 525\ nm$ to detect the green fluorescence of the SYTO9 nucleic acid stain and $560 - 660\ nm$ to detect the chlorophyll autofluorescence of algae. The microscope images showed that an even distribution of algal and bacterial cells was achieved across the filter in a relatively uniform layer. The orthogonal views obtained from a series of z-stack images (with a $2\ \mu m$ step size) confirmed that the vacuum filtration achieved an approximate monolayer of cells.

***Laser marking and gold coating.*** A single hole punch was used to cut out $4 - 6\ mm$ disks from the filter samples. Following this, a Zeiss laser micro-dissection microscope (Zeiss LSM710-NLO housed at the LCI facility of the Karolinska Institute, Stockholm) was used to laser-mark the filter samples and to image the autofluorescence of the algal chlorophyll using the FITC and Rhodamine filter sets. The laser markings could be seen with the camera of the SIMS instrument, and so the SIMS measurements could be matched to chosen sample areas corresponding to particular algal cells in the fluorescence images. After laser-marking the filter samples, they were placed on a conductive sticky tape and mounted onto a glass disk to be placed in the sample holder of the SIMS instrument. The samples were then sputter coated with gold at the NordSIM facility to ensure conductivity of the sample.

#### *Data analysis using WinImage*

The WinImage2 software (Cameca) was used to calculate the isotope ratio $R = {}^{13}C/{}^{12}C$ for single cells of algae and bacteria from SIMS measurements. For bacterial cells, the elliptical tool was used to select regions of interest (ROIs) in the ${}^{12}C^{14}N^{-}$. The isotope ratio for each cell was calculated by taking the mean value for the $100$ scans of SIMS measurements, from which the atomic fraction of ${}^{13}C$, i.e. $f = {}^{13}C/({}^{13}C + {}^{12}C)$ was calculated using

$$f = \frac{R}{1+R}. \qquad (3)$$

During the SIMS analysis there were a few fields of view that contained region(s) of a size comparable to bacterial cells and with a relatively high atomic fraction of ${}^{13}C$. Points were considered outliers and not included in the calculation of the mean if they had an atomic fraction of $^{13}C$ greater than $f_{\max} = p_2 + 4 \cdot (p_2 - p_1)$, where $p_1$ and $p_2$ are the $25th$ and $75th$ percentile respectively. These regions might correspond to bacteria with a relatively high DIC uptake rate or could be the result of an experimental artefact, for example cross-contamination between samples. The rare occurrence of cross-contamination can occur during sample preparation or inside the SIMS instrument. Sputtering with the primary ion beam could cause material from one sample to be deposited on a neighbouring sample, or due to the close proximity of the first lens to the sample surface, material from one sample can land on the mechanical structure of the lens and subsequently be re-deposited onto a different sample (2,3). Future work could benefit from further consideration of this potential for cross-contamination, taking care to not arrange samples too tightly and to minimise the swapping between samples in the run sequence of SIMS analysis.

For algae, the SIMS results for highly labelled cells showed an inhomogeneous distribution of the different carbon isotopes. Therefore, in order to select algal cells in a way that was not biased towards a particular carbon isotope, a linear combination image was created by a simple addition of the two isotope counts $(1 \cdot {}^{12}C^{14}N + 1 \cdot {}^{13}C^{14}N)$, which gives the total distribution of carbon across the area scanned. By comparing this with the fluorescence images, the ROIs corresponding to algal cells were selected. The isotope ratio $R$ was calculated by taking the mean for the $100$ scans of each measurement, from which the atomic fraction of $^{13}C$ was calculated using equation (7). For the preliminary experiment only one measurement of $100$ scans was completed, whereas for the final experiment $2-8$ repeated measurements for each algal cell was obtained.

#### *Depth analysis*

The isotope content can be heterogeneously distributed within the cell and therefore a depth analysis was performed by obtaining repeated measurements of the same cells. SIMS is a destructive technique, meaning that through the process of measurement, as the primary ion beam scans across the sample, the cellular biomass is gradually degraded. For algal cells, the first measurement resulted in only partial degradation of the algal biomass. For three repeated measurements of the atomic fraction $f$ of $^{13}C$ for the same algal cells, the difference between the third and first measurements ($\Delta f = f_3 - f_1$) was calculated relative to the mean $\left(\bar{f} = (f_1 + f_2 + f_3)/3\right)$. The results showed that for the majority of algal cells $f$ either increased or decreased for repeated measurements (Figure S2A), suggesting that the $^{13}C$-enrichment of algal cells was not homogeneous. In order to obtain a measurement that was representative of the whole cell, the mean of three repeated measurements was taken as the value for $f$ of an individual algal cell (with the exception of two cells for the $6\ h$ sample of the pre-labelling culture of algae, for which only two repeated measurements were taken). For bacterial cells, most of the biomass was degraded after the first measurement (Figure S2B), therefore one measurement was sufficient for analysing the carbon isotope content of bacteria.

#### *Scattering effect for highly labelled algae*

When the SIMS scan area contained a labelled algal cell, the $f$ values for bacterial cells in that area were both higher and more variable (Figure S2C). Proximity of algal and bacterial cells on the filter did not necessarily mean physical proximity during growth in the co-culture. Therefore the increase in $f$ for bacteria close to algae on the filter was not simply due to preferential access to DOC exudate. As the caesium ion beam is scanned across the sample, the cellular material is sputtered away to produce secondary ions. However, some of the algal biomass may not be captured as secondary ions and could instead be scattered on the filter in the region around the algal cell. This could explain the observed increase in the mean and standard deviation of the atomic fraction of $^{13}C$ for bacterial cells analysed in the same area as a labelled algal cell. As a result of this observation, only bacteria from scan areas that did not contain labelled algae were included in the analysis described in this work.

##### *Dilution effect - comparing bulk and single cell measurements*

The sample preparation for SIMS analysis introduced unlabelled carbon into the cells during chemical fixation and nucleic acid staining, therefore the atomic fraction of $^{13}C$ was diluted. As established by (4), the relationship between the atomic fraction measured by SIMS $f_{SIMS}$ and the atomic fraction for the sample before chemical fixation and staining $f$ is

$$f = f_{SIMS} + D\ (f_{SIMS} - f_{ch}), \quad (4)$$

where $D$ is the dilution factor and $f_{ch}$ is the atomic fraction of $^{13}C$ in the chemical fixative, and the nucleic acid stain, which were both assumed to be at natural abundance, i.e. $f_{ch} = 0.0108$. The samples for IRMS analysis, which was used for bulk analysis of the carbon isotope content, did not undergo any chemical fixation or staining. Therefore, the IRMS results were assumed to give the true, undiluted value for $f$. To estimate the dilution factor $D$, equation (8) was fitted to the SIMS and IRMS data using $f = f_{IRMS}$ (Figure S2D and Table S3). For bacteria, the fit was carried out using data from only the axenic cultures. In subsequent analysis, to estimate the undiluted atomic fraction of $^{13}C$, the SIMS results were *dilution-corrected* using equation (8) and the dilution factors $D_a = 0.04$ for algal cells and $D_b = 1.29$ for bacterial cells (Table S3). The dilution factor is higher for bacteria than for algae. This is likely to be because the bacterial cells were approximately 10 times smaller than the algal cells and therefore had a greater surface area to volume ratio, which could account for a greater uptake of the chemical fixative and nucleic acid stain.

#### Nutrient-explicit model of the algal-bacterial co-culture

A mechanistic model for the co-culture between a $B_{12}$-dependent alga and a $B_{12}$-producing bacterium, shown schematically in Figure 1, was formulated to capture the population growth, nutrient exchange and isotope labelling kinetics. The model extends a nutrient-explicit model developed by (5) by including more details of the carbon kinetics in order to connect the model to observable variables in isotope labelling experiments. The co-culture model assumes a well-mixed co-culture and provides a nutrient explicit description of an algal-bacterial mutualism that does not rely on detailed metabolic fluxes. The model defines an algal population growth that depends on the external $B_{12}$ concentration $v$ and a bacterial population growth that depends on the DOC concentration, modelled as an effective single carbon source $c_o$, such that

$$\frac{da}{dt} = \mu_a\, a\left(1-\frac{a}{K_a}\right)\left(\frac{v}{K_v+v}\right) \quad \text{and} \quad \frac{db}{dt} = \mu_b\, b\left(1-\frac{b}{K_b}\right)\left(\frac{c_o}{K_c+c_o}\right), \tag{5}$$

with $a$ and $b$ the algal and bacterial cell densities respectively, $\mu_a$ and $\mu_b$ the maximum growth rates, $K_a$ and $K_b$ the carrying capacities, and $K_v$ and $K_c$ the half-saturation concentrations. The internal $B_{12}$ recycling dynamics for algae are neglected and the total $B_{12}$ uptake rate for the whole algal population is given by

$$r_v = \frac{\mu_a\, a}{Y_{a,v}}\left(\frac{v}{K_v+v}\right), \tag{6}$$

with $Y_{a,v}$ the $B_{12}$ yield for algae in the exponential growth phase, i.e. $a \ll K_a$. It is assumed that there is a constant $B_{12}$ production rate per bacterial cell $p_v$, meaning that

$$\frac{dv}{dt} = p_v\, b - r_v. \tag{7}$$

The carbon biomass concentrations for algae and bacteria are given by

$$c_a = \frac{a}{Y_{a,c}} \quad \text{and} \quad c_b = \frac{b}{Y_{b,c}} \tag{8}$$

respectively, with $Y_{a,c}$ and $Y_{b,c}$ the carbon yield parameters, which are assumed to be constant. For simplicity, the model does not consider the carbon concentrating mechanism of *C. reinhardtii* explicitly, instead carbon dioxide and bicarbonate are considered as one entity (i.e. DIC) and photosynthetic assimilation of DIC corresponds to the uptake of both forms of inorganic carbon. Total carbon must be conserved and therefore the rate of photosynthetic carbon assimilation is given by

$$r_p = \frac{\dot{a}}{Y_{a,c}} + r_e, \tag{9}$$

with $\dot{a}/Y_{a,c}$ the algal carbon biomass growth rate and $r_e$ the total rate of DOC exudation by the whole algal population, which is assumed to be linearly dependent on the algal cell density. One further consideration is that there might be an unequal contribution to DOC exudation from different components of the algal biomass. The algal carbon biomass is split into two internal components: the *photosynthetically-active* carbon $c_{a,p}$ and the *stored* carbon $c_{a,s}$. The *stored* carbon corresponds to carbon stored in the form of molecules like starch, but also the carbon used for growth and as a building block for the cellular architecture and machinery. We define the fraction of carbon '*stored'* by algae as

$$\phi_s = \frac{c_{a,s}}{c_a}, \tag{10}$$

from which a *rate of storage* can be defined as

$$r_s = \dot{c}_{a,s} = \frac{\phi_s\, \dot{a}}{Y_{a,c}}\,. \tag{11}$$

The total DOC production rate for the whole algal population is given by

$$r_e = (1 - \phi_s)\, p_c\, a, \tag{12}$$

with $p_c$ assumed to be a constant that can be interpreted as a measure of the rate of DOC production per unit of *photosynthetically-active* algal biomass.

Heterotrophic bacteria must respire to produce the ATP required to drive cellular metabolism, therefore a significant fraction of the carbon consumed by bacteria will be transformed into carbon dioxide through respiration. The bacterial growth efficiency (BGE) is defined as

$$\eta' = \frac{\dot{c}_b}{\dot{c}_b + r_r}, \tag{13}$$

with $\dot{c}_b$ the bacterial carbon biomass growth rate and $r_r$ the respiration rate. In order for the carbon fluxes to remain balanced, while also maintaining an active carbon turnover at carrying capacity, $\eta'$ decreases to zero as the bacterial cell density increases to carrying capacity. Therefore

$$\eta' = \eta\, \left(1 - \frac{b}{K_b}\right), \tag{14}$$

with $\eta$ the maximum BGE, i.e. the growth efficiency for the exponential growth phase when $b \ll K_b$. The total rate of carbon uptake by bacteria is given by $r_u = \dot{c}_b + r_r$ and so from equations (9), (12), (17) and (18)

$$r_u = \frac{\mu_b\, b}{\eta\, Y_{b,c}} \left(\frac{c_o}{K_c + c_o}\right) \tag{15}$$

and the total bacterial respiration rate is therefore given by

$$r_r = \left(1 - \eta \left(1 - \frac{b}{K_b}\right)\right) r_u. \tag{16}$$

Heterotrophic bacteria are able to assimilate inorganic carbon in addition to organic carbon. The model incorporates this observation by including a DIC uptake parameter $X$, defined as

$$X = \frac{r_u^{DIC}}{r_u}, \tag{17}$$

with $r_u^{DIC}$ the DIC uptake rate and $r_u$ the total carbon uptake rate. Taking all these different contributions to the carbon kinetics into account, the model defines the rate of change of the DOC concentration as

$$\frac{dc_o}{dt} = r_e - (1 - X)r_u \tag{18}$$

and the rate of change of the DIC concentration as

$$\frac{dc_i}{dt} = r_r - X\, r_u - r_p. \tag{19}$$

In summary, the mechanistic model describes algal growth dependent on $B_{12}$ produced by bacteria, with photosynthetic uptake of DIC accounting for the algal carbon biomass growth and DOC exudation. The bacterial growth is dependent on the DOC produced by algae, respiration produces carbon dioxide (DIC) and provides the bacteria with the energy they require to grow. In addition to DOC uptake, bacteria are also able to assimilate DIC through metabolic carboxylation reactions (6,7).

#### *Deriving atomic fractions from the nutrient-explicit model*

The atomic fraction of $^{13}C$ is defined as the concentration of $^{13}C$ relative to the total carbon concentration, i.e. $f = {}^{13}C/({}^{13}C + {}^{12}C)$. Each of the different carbon components of the model (bacterial carbon, DOC, DIC etc.) can be considered as a separate carbon pool. For a general case, we consider the $nth$ carbon pool $c_n$ with $r_{gain}$, the rate of carbon coming from the $c_{n-1}$ pool and $r_{loss}$, the rate of carbon going to $c_{n+1}$. This can be summarised as

$$c_{n-1} \xrightarrow{r_{gain}} c_n \xrightarrow{r_{loss}} c_{n+1}.$$

For example, when considering the DOC, $c_n = c_o$, then $c_{n-1} = c_{a,p}$ and $c_{n+1} = c_b$ (the photosynthetic component of algal carbon and the bacterial carbon respectively), $r_{gain} = r_e$ (the rate of DOC exudation by algae) and $r_{loss} = X\, r_u$ (the rate of DOC uptake by bacteria), giving

$$c_{a,p} \xrightarrow{r_e} c_o \xrightarrow{X\, r_u} c_b.$$

The rate of change of the total carbon concentration for the $nth$ carbon pool is

$$\frac{dc_n}{dt} = r_{gain} - r_{loss} \qquad (20)$$

and the rate of change of the $^{13}C$ concentration for the $nth$ carbon pool is

$$\frac{dc_n^{13}}{dt} = f_{n-1}\, r_{gain} - f_n\, r_{loss}, \qquad (21)$$

where $f_n$ is the atomic fraction of $^{13}C$ in the $nth$ carbon pool and $f_{n-1}$ is the atomic fraction of $^{13}C$ in the $(n-1)th$ carbon pool. From equations (24) and (25), the rate of change of the atomic fraction of $^{13}C$ in the $nth$ carbon pool can be derived using the quotient rule, giving

$$\frac{df_n}{dt} = (f_{n-1} - f_n)\frac{r_{gain}}{c_n}. \qquad (22)$$

This general result for the isotope labelling dynamics of a carbon pool assumes that isotopic fractionation is negligible, which corresponds to the assumption that the difference between the nutrient rates for the different carbon isotopes is negligible compared to the overall labelling rates. This assumption leads to the conclusion that the rate of loss to the $c_{n+1}$ carbon pool for the $^{13}C$ and $^{12}C$ isotopes are equal, meaning that the rate of isotope labelling is explicitly independent of $r_{loss}$. However, $r_{loss}$ matters implicitly when comparing the growth and isotope labelling rates (i.e. substituting $r_{gain}$ in equation (26) with $r_{gain} = \dot{c}_n + r_{loss}$ from equation (24) gives $\dot{f}_n = (f_{n-1} - f_n)(\dot{c}_n + r_{loss})/c_n$). If $r_{loss}$ is neglected, then $\dot{f}_n$ would overestimate $\dot{c}_n$, i.e. the labelling rate would overestimate the growth rate.

The algal carbon biomass of the model has two different internal carbon components, meaning that the carbon isotope labelling dynamics for algae does not follow the general case discussed above. In the model, the DOC produced by algae comes from only the photosynthetically active component of the algal biomass and therefore when $f_{a,s}$ is not equal to $f_{a,p}$, the rate of loss for $^{13}C$ and $^{12}C$ from the total algal biomass are not equal. The rate of algal carbon biomass growth is

$$\frac{dc_a}{dt} = r_p - r_e = \frac{\dot{a}}{Y_{a,c}} \qquad (23)$$

and the rate of change of the $^{13}C$ concentration for the algal carbon pool is

$$\frac{dc_a^{13}}{dt} = f_i\, r_p - f_{a,p}\, r_e, \tag{24}$$

from which the differential equation for the atomic fraction of $^{13}C$ in algae

$$\frac{df_a}{dt} = (f_i - f_a)\mu_a\left(1 - \frac{a}{K_a}\right)\left(\frac{v}{K_v+v}\right) + \left(f_i - f_{a,p}\right)(1-\phi_s)\, p_c\, Y_{a,c} \tag{25}$$

is obtained. In contrast to the general case outlined above, the rate of change of the atomic fraction of $^{13}C$ in algae includes a term for the rate of carbon loss.

Taking into consideration the $^{13}C$ isotope labelling dynamics in the general case for the $nth$ carbon pool and in the specific case for algae, the rate of change for the atomic fractions of $^{13}C$ in the co-culture are obtained, giving

$$\frac{df_a}{dt} = (f_i - f_a)\,\mu_a\left(1 - \frac{a}{K_a}\right)\left(\frac{v}{K_v+v}\right) + \left(f_i - f_{a,p}\right)(1-\phi_s)\, p_c\, Y_{a,c}, \tag{26}$$

$$\frac{df_{a,p}}{dt} = \left(f_i - f_{a,p}\right)\left[\left(1 - \frac{a}{K_a}\right)\left(\frac{\mu_a}{1-\phi_s}\right)\left(\frac{v}{K_v+v}\right) + p_c\, Y_{a,c}\right], \tag{27}$$

$$\frac{df_{a,s}}{dt} = \left(f_{a,p} - f_{a,s}\right)\mu_a\left(1 - \frac{a}{K_a}\right)\left(\frac{v}{K_v+v}\right), \tag{28}$$

$$\frac{df_b}{dt} = \left(X\, f_i + (1-X) f_o - f_b\right)\frac{\mu_b}{\eta}\left(\frac{c_o}{K_c+c_o}\right), \tag{29}$$

$$\frac{df_o}{dt} = \left(f_{a,p} - f_o\right)(1-\phi_s)\frac{p_c\, a}{c_o}, \tag{30}$$

$$\frac{df_i}{dt} = (f_b - f_i)\left(1 - \eta\left(1 - \frac{b}{K_b}\right)\right)\frac{\mu_b\, b}{Y_{b,c}\,\eta\, c_i}\left(\frac{c_o}{K_c+c_o}\right), \tag{31}$$

with $f_a$, $f_{a,p}$, $f_{a,s}$, $f_b$, $f_o$ and $f_i$ the atomic fractions of $^{13}C$ in the total algal carbon biomass, *photosynthetically-active* algal carbon, *stored* algal carbon, bacterial carbon, DOC and DIC respectively. This illustrates how a nutrient-explicit model can be used to derive equations for isotope labelling dynamics, allowing the model to make predictions that can be experimentally tested.

#### *Non-dimensional model*

It is instructive to nondimensionalise the co-culture model in order to obtain the minimal set of parameters that characterise the general behaviour of the model. The algal and bacterial cell densities were nondimensionalised using their carrying capacities, that is $\hat{a} = a/K_a$ and $\hat{b} = b/K_b$ respectively.

The $B_{12}$ concentration was rescaled using the half-saturation concentration for algal growth, that is $\hat{v} = v/K_v$. All the carbon concentrations were rescaled using the half-saturation concentration for bacterial growth, that is $\hat{c} = c/K_c$. To nondimensionalise time the bacterial maximum growth rate was used, that is $\hat{t} = t\,\mu_b$. See Table S6 for the definitions of the non-dimensional parameters $\varepsilon$, $k_{a,v}$, $k_{a,c}$, $k_{b,c}$, $s_v$ and $s_c$. From these definitions the non-dimensional ODEs are

$$\frac{d\hat{a}}{d\hat{t}} = \varepsilon\,\hat{a}\,(1-\hat{a})\left(\frac{\hat{v}}{1+\hat{v}}\right), \qquad \frac{d\hat{b}}{d\hat{t}} = \hat{b}\,(1-\hat{b})\left(\frac{\hat{c}_o}{1+\hat{c}_o}\right), \tag{32}$$

$$\frac{d\hat{c}_o}{d\hat{t}} = r_e - (1-X)r_u, \qquad \frac{d\hat{c}_i}{d\hat{t}} = r_r - X\,r_u - r_p, \tag{33}$$

$$\frac{d\hat{v}}{d\hat{t}} = \varepsilon\,s_v\,\hat{b} - r_v. \tag{34}$$

The non-dimensional carbon biomass conversion relations are

$$\hat{c}_a = k_{a,c}\,\hat{a}, \qquad \hat{c}_b = k_{b,c}\hat{b}, \tag{35}$$

$$\hat{c}_{a,s} = \phi_s\,\hat{c}_a, \qquad \hat{c}_{a,p} = \hat{c}_a - \hat{c}_{a,s}. \tag{36}$$

The non-dimensional metabolite rates are

$$r_s = \phi_s\,k_{a,c}\frac{d\hat{a}}{d\hat{t}}, \qquad r_e = (1-\phi_s)\,s_c\,\hat{a}, \tag{37}$$

$$r_p = k_{a,c}\frac{d\hat{a}}{d\hat{t}} + r_e, \qquad r_v = \varepsilon\,k_{a,v}\,\hat{a}\left(\frac{\hat{v}}{1+\hat{v}}\right), \tag{38}$$

$$r_u = \frac{k_{b,c}\,\hat{b}}{\eta}\left(\frac{\hat{c}_o}{1+\hat{c}_o}\right), \qquad r_r = \left(1-\eta\left(1-\hat{b}\right)\right)r_u. \tag{39}$$

The non-dimensional ODEs for the atomic fractions are

$$\frac{df_a}{d\hat{t}} = (f_i - f_a)\,\varepsilon\,(1-\hat{a})\left(\frac{\hat{v}}{1+\hat{v}}\right) + \left(f_i - f_{a,p}\right)\frac{(1-\phi_s)\,s_c}{k_{a,c}}, \tag{40}$$

$$\frac{df_{a,p}}{d\hat{t}} = \left(f_i - f_{a,p}\right)\left[\frac{\varepsilon(1-\hat{a})}{(1-\phi_s)}\left(\frac{\hat{v}}{1+\hat{v}}\right) + \frac{s_c}{k_{a,c}}\right], \tag{41}$$

$$\frac{df_{a,s}}{d\hat{t}} = \left(f_{a,p} - f_{a,s}\right)\varepsilon\,(1-\hat{a})\left(\frac{\hat{v}}{1+\hat{v}}\right), \tag{42}$$

$$\frac{df_b}{d\hat{t}} = (X\,f_i + (1-X)f_o - f_b)\frac{1}{\eta}\left(\frac{\hat{c}_o}{1+\hat{c}_o}\right), \tag{43}$$

$$\frac{df_o}{d\hat{t}} = \left(f_{a,p} - f_o\right)\frac{(1-\phi_s)\,s_c\,\hat{a}}{\hat{c}_o}, \tag{44}$$

$$\frac{df_i}{d\hat{t}} = (f_b - f_i)\frac{\left(1-\eta\left(1-\hat{b}\right)\right)k_{b,c}\,\hat{b}}{\eta\,\hat{c}_i}\left(\frac{\hat{c}_o}{1+\hat{c}_o}\right). \tag{45}$$

#### *Fixed point*

In order for the model to describe a real system the fixed point must have positive values. Therefore, the equations defining the fixed point can be used to derive parameter constraints, which ensure that over time the model variables tend towards positive values.

For the co-culture model developed here, a non-zero fixed point exists where the algal cell density, bacterial cell density, DOC concentration and vitamin $B_{12}$ concentration are all constant. The fixed point for the non-dimensional model is obtained by setting $\frac{d\hat{a}}{dt} = \frac{d\hat{b}}{dt} = \frac{d\hat{c}_o}{dt} = \frac{d\hat{v}}{dt} = 0$, giving

$$\hat{a}^* = 1, \quad (46)$$

$$\hat{b}^* = 1, \quad (47)$$

$$r_e = (1 - X) r_u \;\rightarrow\; \hat{c}_o^* = \frac{(1-\phi_s)s_c}{(1-X)\frac{k_{b,c}}{\eta} - (1-\phi_s)s_c}, \quad (48)$$

$$r_v = \varepsilon\, s_v\, \hat{b}^* \;\rightarrow\; \hat{v}^* = \frac{s_v}{k_{a,v} - s_v}. \quad (49)$$

At this fixed point, the algal and bacterial populations have reached carrying capacity, the rate of DOC production by algae is equal to the rate of DOC uptake by bacteria and the rate of $B_{12}$ production by bacteria is equal to the rate of $B_{12}$ uptake by algae. It is not relevant to consider the case where the DIC concentration is constant, because the model assumes that DIC is in excess and does not affect the rate of algal or bacterial growth.

In order for this fixed point to exist at positive values of $\hat{c}_o^*$ and $\hat{v}^*$, the parameters of the model must satisfy the inequality constraints

$$(1 - X)\frac{k_{b,c}}{\eta} - (1 - \phi_s)\, s_c > 0, \quad (50)$$

$$k_{a,v} - s_v > 0, \quad (51)$$

The isotope labelling dynamics reach a fixed point when all the atomic fractions of $^{13}C$ are equal (i.e. $f_i^* = f_o^* = f_a^* = f_{a,p}^* = f_{a,s}^* = f_b^* = f^*$). This fixed point is defined as

$$f^* = \frac{f_i(0)\, c_i(0) + f_o(0)\, c_o(0) + f_a(0)\, c_a(0) + f_b(0)\, c_b(0)}{c_i(0) + c_o(0) + c_a(0) + c_b(0)}, \quad (52)$$

which can be intuitively understood as simply the weighted average of the initial atomic fractions of $^{13}C$ present in the system. This fixed point depends on the initial conditions, since it depends on the total amount of $^{13}C$ in the co-culture system.

Using the extended co-culture model equations in their non-dimensional form, the Jacobian matrix

$$J = \begin{pmatrix} \frac{\varepsilon\,(1-2\hat{a})\,\hat{v}}{1+\hat{v}} & 0 & 0 & \frac{\varepsilon\,\hat{a}\,(1-\hat{a})}{(1+\hat{v})^2} \\ 0 & \frac{(1-2\hat{b})\,\hat{c}_o}{1+\hat{c}_o} & \frac{\hat{b}\,(1-\hat{b})}{(1+\hat{c}_o)^2} & 0 \\ s_c(1-\phi_s) & -\frac{(1-X)\,k_{b,c}\,\hat{c}_o}{\eta\,(1+\hat{c}_o)} & -\frac{(1-X)\,k_{b,c}\,\hat{b}}{\eta\,(1+\hat{c}_o)^2} & 0 \\ -\frac{\varepsilon\,k_{a,v}\,\hat{v}}{1+\hat{v}} & \varepsilon\,s_v & 0 & -\frac{\varepsilon\,k_{a,v}\,\hat{a}}{(1+\hat{v})^2} \end{pmatrix} \quad (53)$$

was obtained for the ordinary differential equations describing the rate of change of the algal cell density, bacterial cell density, DOC concentration and vitamin $B_{12}$ concentration. The atomic fraction of $^{13}C$ is not included in this analysis because the fixed point $f^*$ in equation (56) and the fixed point for $\hat{a}^*$, $\hat{b}^*$, $\hat{c}_o{}^*$ and $\hat{v}^*$ defined in equations (50)-(53) are independent. In order to determine the stability of the fixed point associated with the population sizes and nutrient concentrations, the Jacobian matrix was evaluated at the fixed point ($\hat{a}^*$, $\hat{b}^*$, $\hat{c}_o{}^*$, $\hat{v}^*$), giving

$$J^* = \begin{pmatrix} -x_1 & 0 & 0 & 0 \\ 0 & -x_2 & 0 & 0 \\ y_1 & -y_1 & -x_3 & 0 \\ -y_2 & y_2 & 0 & -x_4 \end{pmatrix}, \quad (54)$$

with

$$x_1 = \frac{\varepsilon\,s_v}{k_{a,v}},$$

$$x_2 = \frac{(1-\phi_s)s_c}{(1-X)k_{b,c}/\eta},$$

$$x_3 = \frac{(1-X)k_{b,c}}{\eta}\left[1 - \frac{(1-\phi_s)s_c}{(1-X)k_{b,c}/\eta}\right]^2,$$

$$x_4 = \varepsilon\,k_{a,v}\left[1 - \frac{s_v}{k_{a,v}}\right]^2,$$

$$y_1 = s_c(1-\phi_s),$$

$$y_2 = \varepsilon\,s_v.$$

The four eigenvalues of $J^*$ are

$$\lambda = -x_1,\ -x_2,\ -x_3 \text{ or } -x_4, \qquad (55)$$

which are all negative because $x_1$, $x_2$, $x_3$ and $x_4$ are strictly positive (equations (58)). Therefore the fixed point is asymptotically stable, meaning that any small perturbation will converge back to the fixed point (8).

#### Estimating model parameters and solving the model equations

To reduce the number of free parameters, the majority of parameter values were constrained to match values obtained independently from axenic cultures and additional co-culture experiments. The majority of the model parameters were determined using a simplified version of the co-culture model (i.e. with $\phi_s = 0$, $\eta' = 1$ and $X = 0$) to run a global fit of three independent co-culture experiments, which measured colony forming units, particle counts and $B_{12}$ concentrations for a co-culture between *C. reinhardtii metE7* and *M. loti*, see below for details. The algal and bacterial carbon yields were estimated from dry mass measurements and IRMS analysis, see above for details. The remaining parameters were obtained from fitting the model to the stable isotope experiments in this work, see below for details. The full set of model parameters and initial conditions for *C. reinhardtii metE7* and *M. loti* grown both axenically and in co-culture are given in Table S6. Table S7 defines the culture specific parameters and initial conditions for the four axenic cultures of bacteria grown with different concentrations of glycerol. Table S8 compares the results of two parameter optimisations, one with $f_o(0) = 0.64$ and the other with $f_o(0)$ included as a free initial condition. The Matlab ordinary differential equation solver *ode45* was used to numerically solve the model equations.

##### *Parameter optimisation for a simplified co-culture model*

Several of the model parameters were estimated for a co-culture between *C. reinhardtii metE7* and *M. loti* by a global fit of a simplified co-culture model (i.e. with $\phi_s = 0$, $\eta' = 1$ and $X = 0$) to experimental results for three independent co-culture experiments using a basin-hopping algorithm (Figure S5). The co-cultures were grown in Tris-minimal media, $25°C$, $16{:}8\ h$ light:dark cycle and for

$42$, $16$ and $24$ days for experiments $1$, $2$ and $3$ respectively. All three experiments measured colony forming units of *M. loti* and total vitamin $B_{12}$ concentration as determined by bioassy. For experiment $1$ colony forming units of *C. reinhardtii metE7* were used for the fit, whereas for experiments $2$ and $3$ algal counts in terms of particles $> 3\ \mu m$ on the Coulter counter were used. Experiments $1$, $2$ and $3$ include $8$, $4$ and $5$ replicates respectively.

#### *Estimating $K_b$ for axenic bacteria*

To estimate the carrying capacity $K_b$ for axenic cultures of *M. loti*, the logistic growth equation $b = K_b/(1 + M\ e^{-r\ t})$, with positive constant $M$ and $r$, was fit to data taken from (9) for *M. loti* grown axenically with $0.1\ \%$ glycerol. The result is given in Figure S6.

#### *Parameter optimisations using SIMS results*

Parameter optimisations were performed by fitting the nutrient-explicit co-culture model defined above to the population growth and SIMS $^{13}C$-enrichment results. Growth was measured using viable counts and the atomic fractions of $^{13}C$ used were the mean values of the dilution-corrected, single cell measurements obtained using SIMS for each time-point. The Matlab ordinary differential equation solver *ode45* was used to numerically solve the model equations. The parameter optimisations were performed as a global search of the parameter space in order to obtain the best estimate for the set of parameters that minimise the deviation of the model from experiment and that satisfy the boundary conditions (Table S5) and inequality constraints (equations (54) and (55)). Global parameter optimisations were performed using the *GlobalSearch* and *createOptimProblem* functions in Matlab's global optimisation toolbox, with *fmincon* as the solver for each minimisation. All default settings were used except for the *StartPointsToRun* property of the *GlobalSearch* function, which was selected to run with the *bounds-ineqs* option, meaning that all starting points for the minimisations had to lie within the boundary conditions and satisfy the inequality constraints. In order to minimise the number of free parameters, the parameters obtained for the fit of a simplified co-culture model (as outlined above) were used in the parameter optimisations.

#### *Parameter optimisation for the pre-labelling, axenic culture of algae.*

From the non-dimensional co-culture model defined above, an axenic culture of algae can be modelled by setting the initial bacterial concentration to zero (i.e. $b(0) = 0$). Using the experimental data obtained for the pre-labelling, axenic culture of *C. reinhardtii metE7*, the objective function minimised by the parameter optimisation was

$$r^2(a, f_a) = \sum_t \left(\frac{a_{model}(t) - a_{exp}(t)}{a_{exp}(t)}\right)^2 + \left(\frac{f_{a,model}(t) - f_{a,exp}(t)}{f_{a,exp}(t)}\right)^2, \qquad (56)$$

which gives a measure for the deviation of the model from the experiment for both the algal cell density $a$ and atomic fraction of $^{13}C$ for the algal biomass $f_a$. In equation (60) the sum corresponds to the sum over all time-points in the experiment, the subscript $model$ refers to the value obtained from the model and the subscript $exp$ refers to the value measured experimentally.

<u>Free parameters and initial conditions</u>. The free parameters were $s_c$ and $\phi_s$. All other parameter values used were as defined in Supplementary Table S6. For the experiment, it was assumed that initially there was no DOC in the media, the DIC was in excess and the algae were initially unlabelled, therefore $\hat{c}_o(0) = 0$, $\hat{c}_i(0) = 5$, $f_a(0) = 0.0108$ and $f_o(0) = 0.0108$. No reliable measurement for the initial algal cell density was obtained and although the initial $B_{12}$ concentration was $100\ ng\ L^{-1}$, the model for algal growth neglects the internal $B_{12}$ recycling dynamics and so the $B_{12}$ concentrations in the model do not necessarily correspond to the quantitative values of the experiment, therefore the initial algal cell density and $B_{12}$ concentration were kept free. Although the $NaH^{13}CO_3$ used for the stable isotope labelling cultures had $98\ atm\%^{13}C$, due to the equilibria between different forms of inorganic carbon, the actual atomic fraction of $^{13}C$ for the DIC assimilated by the algae was unknown, therefore the initial condition $f_i(0)$ was also kept free.

#### *Parameter optimisation for axenic bacteria*

An axenic culture of bacteria can be modelled using the co-culture model defined above and setting the initial algal cell density to zero (i.e. $a(0) = 0$). A global parameter optimisation was performed for axenic bacteria using the experimental results of four cultures of *M. loti*, each grown with a different concentration of glycerol ($0.1$ %, $0.01$ %, $0.001$ % and no glycerol). The objective function minimised by the global parameter optimisation was

$$r^2(b, f_b) = \sum_{all\ cultures} \sum_t \left(\frac{b_{model}(t) - b_{exp}(t)}{b_{exp}(t)}\right)^2 + \left(\frac{f_{b,model}(t) - f_{b,exp}(t)}{f_{b,exp}(t)}\right)^2, \qquad (57)$$

with the sum over $all\ cultures$ indicating that the aim was to minimise the difference between the model and the experimental results for the bacterial cell density $b$ and the atomic fraction of $^{13}C$for bacteria $f_b$ for all four axenic cultures simultaneously.

***Free parameters***. The model parameters for axenic bacteria were considered as global parameters, with the exception of $\eta$ and $X$ that could have values specific to the different cultures. The maximum growth rate and carbon uptake parameter for bacteria ($K_b$, $\mu_b$ and $k_{b,c}$ respectively) obtained for *M. loti* in co-culture with *C. reinhardtii metE7* might not be the same as for *M. loti* grown in axenic cultures in which bacteria are grown with glycerol as their organic carbon source. Therefore $K_b$ was determined as described above, and $\mu_b$ and $K_c$ were kept as free global parameters. Using $Y_{b,c} = 5 \times 10^{14}\ cells\ molC^{-1}$ obtained from dry mass measurements and IRMS results (as described above), the value for the carbon uptake parameter $k_{b,c}$ was updated throughout the parameter optimisation as $K_c$ changed, according to the parameter definition $k_{b,c} = K_b/(Y_{b,c}\ K_c)$.

***Initial conditions***. It was assumed that in the experiments the DIC was in excess and the atomic fraction of $^{13}C$in the DIC was taken as the estimate obtained from the parameter optimisation for axenic algae, meaning $\hat{c}_i(0) = 5$ and $f_i(0) = 0.65$. Initially, there was no $B_{12}$ in the media, the bacteria were at natural abundance and the glycerol was unlabelled; therefore $\hat{v}(0) = 0$, $f_b(0) = 0.0108$ and $f_o(0) = 0.0108$. The initial DOC concentrations were calculated for $0.1$ %, $0.01$ % and $0.001$ % glycerol concentrations to be $4 \times 10^{-5}$, $4 \times 10^{-6}$ and $4 \times 10^{-7}\ molC\ mL^{-1}$ respectively, using the molar mass of glycerol, $92.09\ g\ mol^{-1}$, and its density, $1.26\ g\ mol^{-1}$. Although it is expected that the axenic culture grown without glycerol had no DOC in the media, the experimental results suggest that there was still a small amount of bacterial growth (Figure 2). In order to account for this observation, $c_o(0)$ for the `no glycerol' culture was kept free, but was constrained to be less than $4 \times 10^{-7}\ molC\ mL^{-1}$ (i.e. $0.001\ \%$ glycerol). No reliable measurement for the initial bacterial cell density was obtained experimentally, therefore $b(0)$ for each culture was also kept free in the global parameter optimisation.

#### *Parameter optimisation for the co-culture*

The objective function of the parameter optimisations for the co-culture was

$$r^2(a, b, f_a, f_b) = \sum_t \left(\frac{a_{model}(t) - a_{exp}(t)}{a_{exp}(t)}\right)^2 + \left(\frac{b_{model}(t) - b_{exp}(t)}{b_{exp}(t)}\right)^2 + \left(\frac{f_{a,model}(t) - f_{a,exp}(t)}{f_{a,exp}(t)}\right)^2 + \left(\frac{f_{b,model}(t) - f_{b,exp}(t)}{f_{b,exp}(t)}\right)^2, \quad (58)$$

which gives a measure for the deviation of the model from the experiment for both the cell densities ($a$ and $b$ for algae and bacteria respectively) and the atomic fractions of $^{13}C$ ($f_a$ and $f_b$ for algae and bacteria respectively).

***Estimating $\phi_s$, $\eta$ and $X$***. The result $\phi_s = 0.9$, obtained from the parameter optimisation for axenic algae, was carried forward for the co-culture model. The results for the axenic bacteria suggested that for a higher initial glycerol concentration, and therefore a higher exponential growth rate, the value for the DIC uptake parameter $X$ increases and the value for the bacterial growth efficiency $\eta$ decreases. These trends were used to estimate $X$ and $\eta$ for the co-culture. For axenic bacteria, the initial glycerol concentration $c_o(0)$ was used to estimate the exponential growth rate $\mu_B = \mu_b\ c_o(0)/(c_o(0) + K_c)$. An exponential growth rate fit for bacteria in the co-culture gave estimates for the initial bacterial cell density $b(0) = 1.2 \times 10^7 \pm 1.5 \times 10^5\ cfu\ mL^{-1}$ and the exponential growth rate $\mu_B = 0.022 \pm 0.005\ h^{-1}$ (Figure S7). A linear fit for $X$ against $\ln(\mu_B)$ (Figure S8A) was used to obtain the estimate $X = 0.015 \pm 0.001$ for bacteria in the co-culture. A linear fit for $\eta$ against $\ln(\mu_B)$ (Figure S8B) was used to obtain the estimate $\eta = 0.51 \pm 0.21$ for bacteria in the co-culture.

***Free parameters and initial conditions***. The majority of the model parameters were fixed with values as defined in Supplementary Table S6, apart from $s_c$, which was included as a free parameter. The initial conditions of the co-culture meant that DIC was in excess and the $B_{12}$ concentration was assumed to be zero (because bacteria were washed thoroughly prior to establishing the co-culture and $B_{12}$ was assumed to have been fully depleted in the pre-labelling culture of algae because it was inoculated with only $100\ ng\ L^{-1}$ $B_{12}$), therefore $\hat{c}_i(0) = 5$ and $\hat{v}(0) = 0$. For the initial atomic fraction of $^{13}C$ in the DIC, the estimate obtained from the parameter optimisation for axenic algae was used, i.e. $f_i(0) = 0.65$. The co-culture was inoculated with pre-labelled algae, therefore the $48\ h$ time-point of the pre-labelling culture was used to estimate the initial atomic fractions of $^{13}C$ in the algae and DOC. Using the model fit results for the axenic algae given in Supplementary Table S6, estimates for the initial conditions $f_a(0) = 0.59$, $f_{a,p}(0) = 0.65$ and $f_o(0) = 0.64$ for the co-culture were obtained. The bacteria started the co-culture at natural abundance and so $f_b(0) = 0.0108$. The initial conditions that remained free during the parameter optimisations were $\hat{a}(0)$, $\hat{b}(0)$ and $\hat{c}_o(0)$, with $f_o(0)$ also included as a free initial condition for fit 2.

### III. Supplementary Results

#### Pre-labelling algae in an axenic culture

The $B_{12}$ dependent *C. reinhardtii metE7* was grown axenically for $48\ h$ in media containing $5\ mM$ $NaH^{13}CO_3$, which provided a $^{13}C$-enriched inorganic carbon source for photosynthesis. For each time-point, SIMS images (Figure S9A) were used to obtain measurements of the atomic fraction of $^{13}C$ in individual algal cells (Figure S9B). The mean $f_a$ was then calculated (Figure S9C), excluding the cells close to natural abundance, which are highlighted in red in S9B, because they were assumed to be inactive and not contribute to the carbon dynamics of the culture. The value for $f_a$ increased throughout the culture, indicating that *C. reinhardtii metE7* used the $^{13}C$-enriched DIC for photosynthesis and growth (Figure S9C-D). The rate of $^{13}C$-enrichment decreased as the culture progressed, with $f_a$ beginning to plateau (Figure S9C). It is likely that the labelling rate decreases when $f_a$ approaches the value of $f_i$, meaning that $f_a$ and $f_i$ reach an equilibrium. Although the $NaH^{13}CO_3$ used had an atomic fraction of $^{13}C$ of $0.98$, due to the equilibria between different forms of DIC and atmospheric carbon dioxide, the actual atomic fraction of $^{13}C$ for the DIC assimilated by the algae is unknown. The model achieved a good fit to the experimental data for the axenic culture of algae (Figure S9 and Table S6) and estimated the initial atomic fraction of $^{13}C$ in the DIC to be $f_i(0) = 0.65$. This value for $f_i(0)$ was used in the parameter optimisations for the axenic cultures of bacteria and the co-culture. In the model, as algae become labelled, the DOC they exude also becomes labelled. Using the parameter optimisation results given in Table S6, it was found that after 48 hours of the pre-labelling culture of algae, $f_a = 0.59$, $f_{a,p} = 0.65$ and $f_o = 0.64$. These values were used as initial conditions for the model fit of the co-culture.

#### Comparing SIMS results of two independent experiments

A preliminary experiment for the SIMS analysis was performed for a pre-labelling culture of axenic algae, a labelled co-culture and an axenic culture of bacteria with $0.1$ % glycerol. The results from the preliminary SIMS experiment show the same trends in the isotope labelling dynamics as those observed for the final SIMS experiment (Figure S10). This illustrates the repeatability of the measurements obtained. The unlabelled control cultures for axenic algae and a co-culture were included in the preliminary experiments and showed the expected result of natural abundance.
